# Structured experience shapes strategy learning and neural dynamics in the medial entorhinal cortex

**DOI:** 10.1101/2025.05.13.653873

**Authors:** John C. Bowler, Dua Azhar, Cambria M Jensen, Hyun-Woo Lee, James G. Heys

## Abstract

Animals can solve new, complex tasks by reusing and adapting what they’ve learned before. This kind of flexibility depends not just on having prior experience, but on how that experience was structured in the first place. The design of early training curriculum is especially important: poorly structured experiences can hinder abstraction and limit generalization, while carefully structured training promotes more flexible and adaptive behavior. Yet, the neural mechanisms supporting this process remain unclear. To investigate how early training shapes learning we first trained recurrent neural networks (RNNs) on variants of an odor-timing task previously used to study complex timing behavior in mice. We then tested the RNN predictions on how previous experience affects generalization using behavioral and electrophysiological recordings in mice trained on the same task using staged training sequences. RNNs and mice trained without well-structured early experience developed rigid strategies and made repeated errors. In contrast, those given more balanced early training were better able to generalize and showed similar neural activity patterns that reflected the task’s underlying temporal structure. Using dynamical systems approaches, we reveal a mechanism for this effect: networks trained with appropriately structured curricula developed distinct dynamical motifs that support the correct abstractions when complexity was increased. Networks that lacked early training or received remedial curricula developed single fixed-point solutions that failed to generalize beyond the training stimuli. Together, these findings demonstrate that it is not just the presence of prior experience, but its structure, that governs how flexible and generalizable knowledge emerges in both biological systems and computational models.

## Introduction

Learning enables animals to construct flexible knowledge structures, allowing them to adapt to new tasks by reconfiguring prior experiences into novel strategies. This compositional nature of learning suggests that the structure of prior experiences—the curriculum—is critical for future problem-solving. Poorly structured curricula, however, can cause animals to become ”stuck,” perseverating on sub-optimal solutions that only partially achieve their goals (Le et al. 2023).

Despite the central role of structured experience in both everyday learning and experimental settings, the neural mechanisms through which the organization of past experience shapes future learning remain poorly understood. Therefore, we asked: how does the structure of past experience shape both the cognitive strategies and the underlying neural circuit dynamics that support flexible task performance? Commonly, past experience is conceptualized as a library of atomic units of learned behaviors, called task primitives, each accomplishing a basic goal. Complex behaviors are thought to arise by compiling these primitives into behavioral programs (Chomsky 2017, Elman 1993, Lee et al. 2024, Randlov & Alstrøm 1998), making learning inherently compositional. In laboratory and real-world settings alike, simpler shaping tasks or experiences often break down complex behaviors into sub-units that can later be assembled—or ”composed”—into strategies capable of achieving broader goals (Bengio et al. 2009, Krueger & Dayan 2009, Narvekar et al. 2020, Skinner 1938). However, it is often unclear how the structure of prior training alters cognition (Le et al. 2023). Moreover, shaping protocols—or the natural structure of experience—inevitably embed assumptions about how tasks should be solved, potentially biasing the strategies that emerge. These challenges are amplified when animals must perform distinct actions in contexts with overlapping features, where no single observation reliably signals the correct behavior (Gagliardi et al. 2024, Keene et al. 2016, Lee & Lee 2013, Plitt & Giocomo 2021).

Strategies to solve complex context-dependent behaviors necessitate the involvement of mnemonic reasoning, in order to link events to the problem space and arrive at reasonable approaches based on experience. Critically, medial temporal lobe (MTL) structures are involved in this kind of context-dependent, episodic learning (Buzsáki & Moser 2013, Eichenbaum 2004, Mishkin 1978, Muller et al. 1987, O’Keefe & Dostrovsky 1971, O’Keefe & Nadel 1978, Scoville & Milner 1957, Squire & Zola-Morgan 1991, Wilson & McNaughton 1994, Wood et al. 1999) and exhibit activity patterns that are correlated with the currently selected cognitive strategy (Bigus et al. 2024, Ferbinteanu & Shapiro 2003, Jadhav et al. 2012, Wikenheiser & Redish 2015, Wood et al. 2000). Within the MTL, the medial entorhinal cortex (MEC) is a highly recurrently connected region (Buetfering et al. 2014, Dhillon & Jones 2000, Kumar et al. 2008, Zutshi et al. 2018) that is most famously known for being a focus of grid cell tuning during navigation-based behaviors

(Hafting et al. 2005), but also has been shown to exhibit context-dependent activity (Boccara et al. 2019, Butler et al. 2019, Fyhn et al. 2007, Kitamura et al. 2015) as well as correlates of interval timing (Heys & Dombeck 2018, Heys et al. 2020) and is necessary for complex and context-dependent timing behaviors (Bigus et al. 2024, Heys & Dombeck 2018). Theories persist about how a recurrently connected region such as MEC might generate both spatially and temporally correlated activity based on task demands through attractor dynamics (Issa et al. 2020, Yoon et al. 2013); however, a key question remains unanswered: what mechanisms permit network activity to traverse context specific trajectories? In the MEC, the linking of context-dependent behavior with recurrently generated attractor dynamics must rely on previous experience in order to result in observable neural correlates of cognitive strategy. MEC is therefore a brain region that should exhibit experience-dependent changes in neural activity that could bias strategy learning in the future.

Notably, it has also been shown *in-silico* that pre-training steps similar to those used to train animals results in dynamical systems that adopt similar strategies to those utilized by animals when solving complex problems (Bengio et al. 2009, Hocker et al. 2024, Ito et al. 2022). Recurrent Neural Networks (RNNs) can also be trained to do relatively complex timing comparison tasks, recapitulating key features of the neural dynamics observed in animals (Lin et al. 2023) using plausible network connectivity motifs (Zhou et al. 2023). Indeed, RNNs can be trained to learn and reuse task primitives, similar to animals (Duncker et al. 2020, Yang et al. 2019), and perform multitask learning to investigate how multiple task representations can be composed to support different outcomes based on various learning strategies (Driscoll et al. 2024, Márton et al. 2022, Yang et al. 2019). Therefore, modeling learning in artificial networks can give key insights into how strategies evolve across learning and which approaches animals implement to compose elements of past experience. Critically, much prior work relies on task primitives with strictly orthogonal input spaces (Lee et al. 2024) or seeks to understand tasks with an explicit context identifying input (Driscoll et al. 2024, Mante et al. 2013, Yang et al. 2019), which is not the case in most laboratory settings or real world experiences.

To directly address how the structure of prior experience shapes learning, we took a theory-driven approach combining computational modeling and experimental testing. Our study was motivated by prior observations that mice could adopt fundamentally different strategies to solve the same odor-timing task, suggesting that subtle differences in neural circuit mechanisms—shaped by early training and potentially by individual variability—might underlie these divergent solutions (Bigus et al. 2024). To generate mechanistic predictions, we trained recurrent neural networks (RNNs) on variants of the task, using constraints derived from behavioral shaping procedures in mice. This allowed us to test how the structure of early experience influences the emergence of learning dynamics and cognitive strategies. Critically, analysis of RNN dynamics revealed that well-structured pre-training organizes network activity into distinct low-dimensional motifs that form the computational substrate for abstraction and flexible generalization. In contrast, poorly structured training failed to induce these motifs, resulting in rigid, single-solution dynamics and impaired task performance. We then tested these predictions experimentally, using behavioral and electrophysiological recordings from mice trained with matched shaping sequences. Mice receiving well-structured early experience exhibited better generalization and medial temporal lobe activity patterns resembling those observed in the models. Together, these findings demonstrate that it is not just the presence of prior experience, but its structure, that governs how flexible and generalizable knowledge emerges in both biological and artificial neural systems.

## Results

In order to study how dynamical systems learn to perform complex tasks, we developed a Recurrent Neural Network (RNN) implementation, similar to past studies (Mante et al. 2013, Yang et al. 2019, Duncker et al. 2020, Driscoll et al. 2024), that could be trained on a complex cue timing task previously implemented in mice (Bigus et al. 2024). In this way we could fully observe both the network dynamics and architecture to gain key insight into how dynamical systems, such as our model (or the mouse brain), develop task solutions. These networks can additionally provide insight into how prior experience alters the dynamical landscape pushing the system to use optimal (or suboptimal) strategies. To identify the effect of structured early experience, we incorporate a behavioral shaping process, similar to the behavioral shaping steps used when training animals to perform complex tasks (Bigus et al. 2024), that is critical to the networks’ ability to learn the task.

The task is adopted from an odor timing task in which animals observe two consecutive olfactory stimuli, comprised of the same odorant but presented for distinct durations, with a fixed inter-stimulus interval (ISI). To perform the task correctly, trials of non-matching durations are identified and reported by licking at a reward spout. The durations used were either 2s (“short”, S) or 5s (“long”, L) and three trial types were used in the experiments: one “match,” or no-go trial type Short-Short (SS), along with two “non-match,” or go trial types Short-Long and Long-Short (SL, LS). To solve the task, animals must identify a response window following the second cue offset– licking restricted to this window triggers to a water reward delivery. Licking prior to the onset of the second stimulus during non-match trials, or at anytime during a match/SS trial, is considered an error. The task is therefore a modified temporal variant of delayed non-match to sample (tDNMS). Trials are randomized in blocks of 4 with 2 match (SS) and 2 non-match (SL/LS) trials in each block. For mice, trials were initialed with a 250 ms LED flash. For RNNs, the tDNMS task incorporates two inputs, a “start cue” and a “timed cue,” that are one-hot encoded for off vs on and then altered with Gaussian input noise (*µ* = 0, *σ* = 0.15). The networks we train to perform this task have a fully connected recurrent layer of “leaky” units, each receiving the two inputs and sending a feed-forward connection to a single “response” node (Figure 1b). The networks are trained to identify the response windows on “Go” trials by manipulating the value of the output node (fully connected to the recurrent layer with feed-forward, trainable weights) and trained to minimize error between their output and a target function representing the response window (Figure 1c).

**Figure 1.**
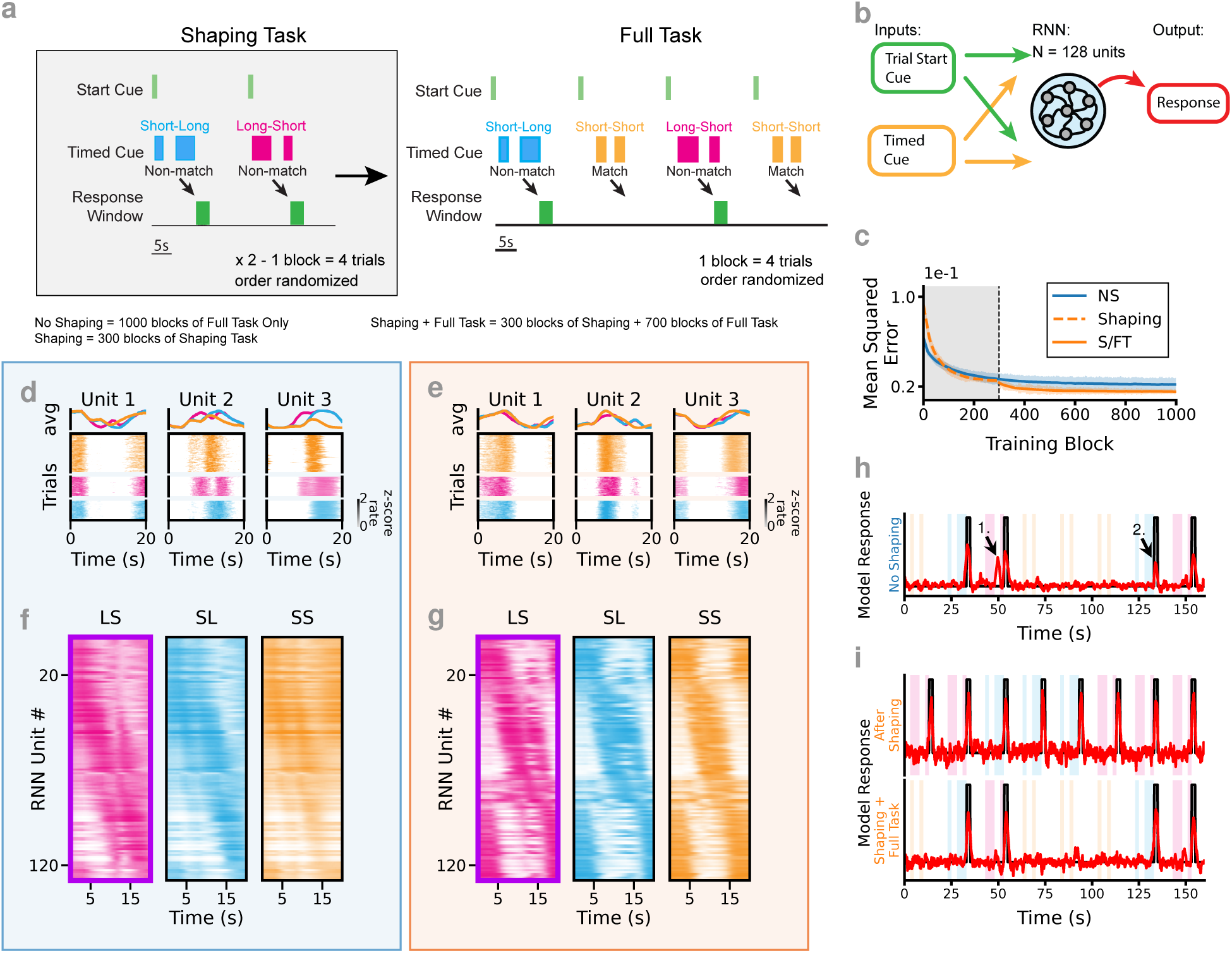
RNNs learn to solve a complex timing task. (**a**) The tDNMS task is comprised of 3 trial types: 2 “Go” trials (Long-Short, LS; Short-Long, SL) and 1 “No-Go” trial type (Short-Short, SS). Trials are presented in blocks of 4 with trial type shuffled within each block. *Left*. During shaping, only the “Go” trials are used and the correct behavior is to withhold responses until the offset of the second cue. *Right*. During the full task the match/“No-Go” trials are introduced, in which response much be withheld entirely. (**b**) RNNs are constructed with a fully connected recurrent layer of ”leaky” units, each receiving feed forward inputs from the trial start cue and a timed cue and sending a feed-forward connection to a single “response” or output node. (**c**) MSE comparison across each block of training for *n* = 12 S/FT RNNs and *n* = 12 NS RNNs mean (line) and standard deviation (shaded region). For the S/FT RNNs the first 300 blocks of training omit the matched short-short/“No-Go” trial type. (**d**) Example unit activity of RNN following training without shaping task. (**e**) Example unit activity on RNN following S/FT training. (**f**) Tuning curves for all 128 units of an NS RNN. (**g**) Tuning curves for all 128 units of an RNN trained using the S/FT protocol. (**h**) Example response for an RNN without shaping (NS). Note the early response on LS trial (1.) and the relatively low amplitude response on SL (2.). (**i**) (*Top*.) Example response of an RNN after training on the shaping task for 300 blocks. (*Bottom*.) Response of the same model after training for an additional 700 blocks on the full task.

Critically, in order for mice to learn the tDNMS task, extensive behavioral shaping is necessary. Shaping takes place in the same apparatus as the full task, familiarizing the animals with the experimental setup while introducing them to the basic rules of the task. During the shaping phase of learning, the match (SS) trial type is withheld from the mice until they reach proficiency on the non-match task (*see Methods*). To isolate how past experience shapes future learning, we augmented the RNNs’ training with an analogous “no match” task prior to exposing the networks to the full task (Figure 1a). Further, we compare networks that have undergone the shaping experience prior to learning the full task (S/FT) with networks that were never trained on the shaping task (NS). We train all networks for 1,000 blocks of 4 trials at which point the behavior of the networks has converged with minimal changes in the mean squared error (MSE) between training blocks (Figure 1c). For S/FT networks, we replace the first 300 blocks with the no match shaping task (subsequently, these networks will be labeled as “Only Shaping” RNNs). Following training, and in agreement with observations of neural activity in mice (Bigus et al. 2024), we observe similar activation patterns in both NS and S/FT RNNs; many individual units exhibit sparsely active temporally tuned fields (Figure 1d,e). At a population level, the activity of individual units in S/FT networks, when compared to units in NS networks, more uniformly tiles the duration of the full task (as observed by sorting the trial averaged activity of each node by LS, Figure 1f,g), however, this difference is somewhat subtle.

We find stereotypical errors in NS RNNs when identifying the response window, particularly early responses following the Long cue on LS trials, along with relatively low amplitude responses to both LS and SL trial types (Figure 1h). This combination makes it difficult to define a cutoff threshold to discriminate between a Go and No-Go trial type (Figure S1a). In contrast, the shaping phase guides these S/FT RNNs to solve the task robustly and without error (Figure 1i, S1a).

### After shaping, a key abstraction about task structure is learned by the RNNs

To visualize how the S/FT RNNs solve the tDNMS task differently from the NS networks, we plot the first 3 principal components (PCs) as calculated on the recurrent layer of the networks. Immediately, it is apparent that NS networks show a very different task representation than the S/FT RNNs (Figure 2a, S2b,a). Particularly the NS networks exhibit a fairly tangled (Russo et al. 2018) pattern of activity without an easily interpretable structure. To directly compare how the networks evolve with learning, state space representations of networks which have had a prior shaping experience (Only Shaping) are projected into the corresponding PCA space of the network after training on the full task. In these networks, the first two PCs indicate that the RNNs traverse a very stereotypical circular subspace that is largely conserved between the 3 trial types (Figure 2a, S2b). The remarkably similar structure of the task representation between trial types is mainly distinguished around the response window in PC3. This deflection also occurs on the SS trial types prior to training on the full task. After incorporating the No-Go trials into training, diminished variance in PCs is observed in the SS trial type corresponding to the network learning to inhibit responses (Figure 2a, *dashed line*). This means only a small change is required to adjust behavior to learn the correct full task behavior. This change is mainly observed in the reduced variance in PC3 around the response window. Additionally, experiencing the shaping task causes RNNs to significantly restrict the variance of the network to the first 3 PCs compared to the NS network (Figure 2b). In this way, the behavior constrains the dynamics to be lower dimensional, and exhibit significantly less tangled trajectories (NS tangling greater than S/FT, *n* = 12 NS and 12 S/FT RNNs, *p* = 0.02, Figure 2c, S2c). Low dimensionality can be enforced in RNNs during training and has been shown to lead to improved generalization during timing tasks (Beiran et al. 2023). Interestingly, subjecting the networks to a correctly designed behavioral shaping task enforces a similar constraint on the resulting fully trained RNN.

**Figure 2.**
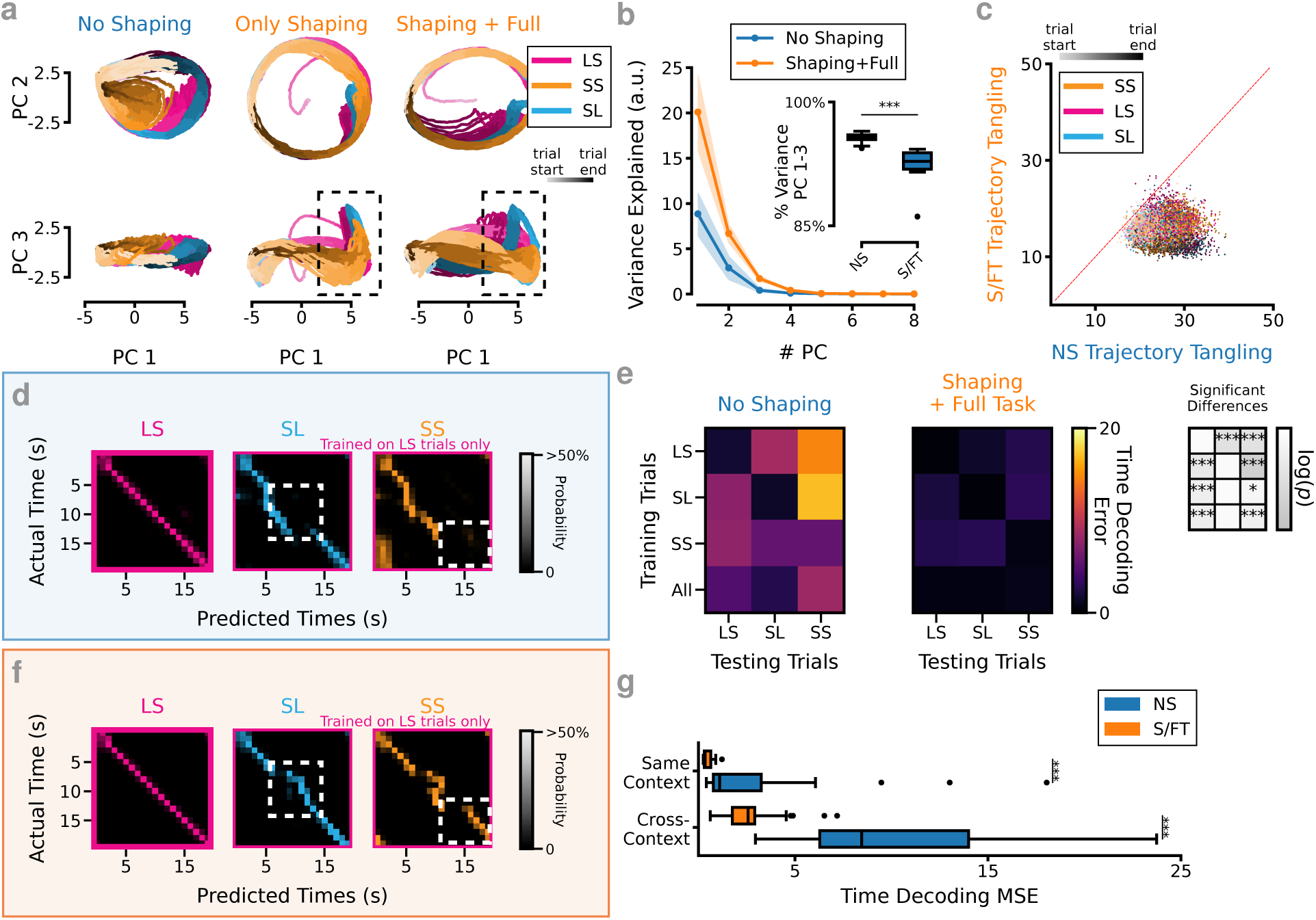
Behavioral shaping results in learning an abstract representation of the trial structure. (**a**) PCA representation of hidden layer state space from RNN without shaping (NS, *left*), after 300 blocks of training on the shaping task (OS, *middle*) and then after training on shaping followed by the full task (S/FT, *right*). *Top*. Plot of PC1 and PC2. *Bottom*. Plot of PC1 and PC3. Dashed boxes indicate key difference between OS and S/FT networks near the response window. OS PCs are projected into the task PCA space of S/FT network. (**b**) S/FT RNNs have more variance explained by fewer PCs, mean and std. of *n* = 12 S/FT RNNs and *n* = 12 NS RNNs, significant effect of shaping experience (*p* = 5.6 *×* 10*^−^*^10^), # PC (*p* = 3.5 *×* 10*^−^*^123^), and the interaction (*p* = 4.5 *×* 10*^−^*^53^), two-way mixed ANOVA. *Inset*. S/FT networks also have a higher percentage of variance explained by the first three PCs, *p* = 2.78 *×* 10*^−^*^4^, two-factor independent t-test. (**c**) NS networks exhibit more tangled trajectories in state space than S/FT networks. Scatter of NS and S/FT networks trajectory tangling metric for matched input across 15 blocks of 4 trials of the full task. (**f**) Example within and across context time decoding for an S/FT RNN. *Left*. LDA based cross-validated decoders trained on LS trials and tested on held out data. *Middle*. The same LS trained decoders tested on SL trials. *Right*. LS trained decoder tested on SS trials. (**e**) *Left/Middle*. Summary of time decoding within and across trials for *n* = 12 NS and *n* = 12 S/FT RNNs. *Right*. *p*-values for NS compared to S/FT results, two-sided independent t-test with Bonferroni correction, *-*p <* 0.05, **-*p <* 0.01, ***-*p <* 0.001. (**d**) Same as f, but for NS RNN. (**g**) S/FT RNNs outperform NS RNNs at decoding trial time for both within- and across-context decoding schemes *n* = 12 NS RNNs, *n* = 12 S/FT RNNs, *p* = 2.3 *×* 10*^−^*^4^ within-context decoding, *p* = 1.3 *×* 10*^−^*^23^ across-context decoding, two-sided independent t-test.

The distinct activity pattern developed during pre-training led us to question: *“What do RNNs learn from the shaping task?”* The evident separation of trial timing (in the first two PCs) from response dynamics (in PC3) led us to question whether the RNNs had learned to time trials independently of the trial type. To test this, we trained a Linear Discriminant Analysis (LDA) based classifier to decode trial time on 10 splits of the unit activity, restricting training either to a specific trial type or all trials. Strikingly, S/FT models could decode the passage of time not only for *within-*context temporal representations (two-sided t-test, *n* = 12 NS, 12 S/FT, *p* = 2.3 × 10^−4^), but also in the cross-context case, performing significantly better than NS RNNs (two-sided t-test, *n* = 12 NS, 12 S, *p* = 1.3 × 10^−23^, Figure 2d-g, S2d,e); cross-context decoding is thought to reflect concept generalization or abstract thinking (Bernardi et al. 2020, Fascianelli et al. 2024). Therefore, the previous shaping experience has not only enforced a lower-dimensional representation of the task into the RNNs, but has also led to learning an abstraction between trial types, corresponding to a generalized concept of elapsed trial time.

### Prior experience persistently alters network connectivity

Training RNNs is most directly the process of learning the connection weights necessary for best task performance. Looking at the connection weights of the RNNs reveals direct evidence for how the shaping experience structurally modifies the networks and provides key insights into differences between the NS and S/FT RNNs. Sorting the connectivity matrices by peak activation time during the tDNMS trials gives us a window to investigate the connectivity patterns and make comparisons between RNNs (Figure 3a,b). Generally, in both the NS and S/FT RNNs, nearby units in the temporal sequence tend to have excitatory connections while cells with more distant peaks show inhibitory connections (Figure 3c). This nearby excitation, surround inhibition pattern is in agreement with attractor network models of how dynamical systems might engage in timing behaviors (Issa et al. 2020, Zhou et al. 2023). While the connection pattern predicts that a bump of activity would propagate through the networks, it is unclear how the observed context-dependent and trial type specific activity patterns are generated from the observed pattern. However, the S/FT network shows alternating blocks of units with highly positive inputs from the timed cue indicating heterogeneity in the tuning curves based on input state in S/FT networks as opposed to NS.

**Figure 3.**
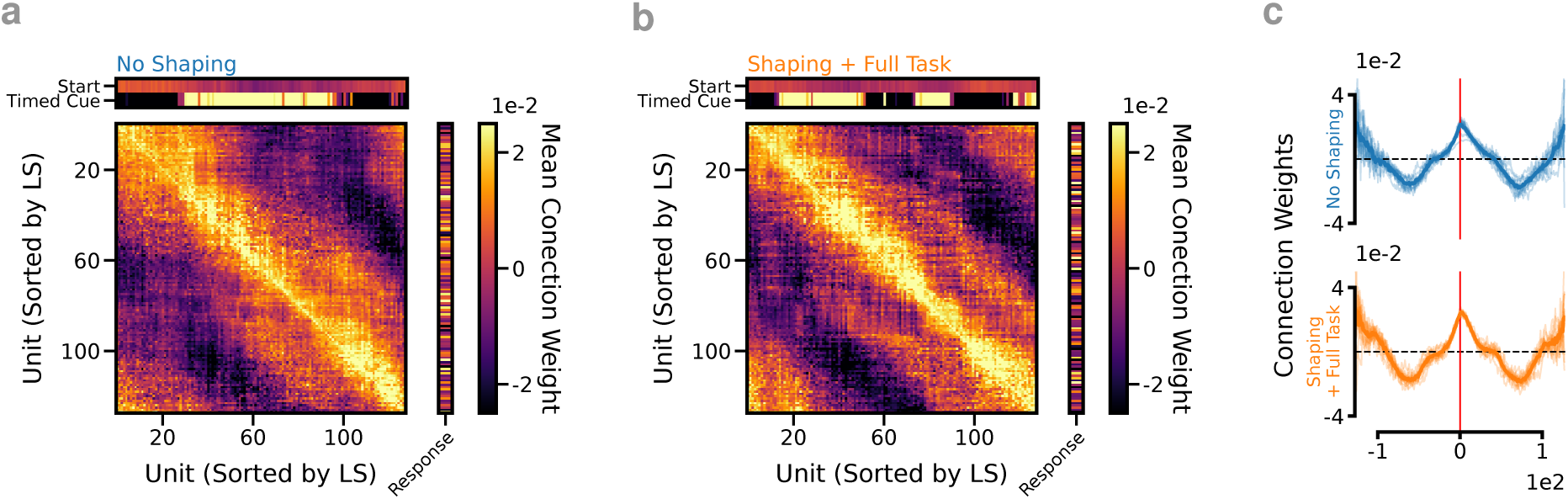
What effect does the shaping task have on the eventual architecture of RNNs after learning? (**a**) Sorting the connectivity matrix from an RNN that was trained without shaping by peak activation on LS trials. Average weight matrices from *n* = 12 No Shaping RNNs. (**b**) Same as a for a network that trained on the shaping task. Average weight matrices from *n* = 12 RNNs trained with Shaping. (**c**) Nearby units in the temporal sequence tend to have excitatory connections while cells with more distant peaks show inhibitory connections.

### Dynamical motifs predict network behavior

Because our analysis of the RNN connection weights resembled standard models of attractor dynamics, we next asked whether the RNNs’ activity patterns during the task could be explained by identifying their underlying dynamical motifs (Driscoll et al. 2024). To do this, we systematically searched for fixed points and slow points within the region of state space occupied by the RNNs during the tDNMS task. Importantly, this analysis is performed after training is complete and the RNN weights are unchanging, therefore the network evolution is fully determined by the current location in this state space and the external input. Focusing on the timed cue (since the start cue was brief and identical across trials), we separated network dynamics into two main ”modes”: Cue On and Cue Off. Using numerical methods (Sussillo & Barak 2013, Golub & Sussillo 2018), we identified fixed points across NS, Only Shaping, and S/FT networks. Additionally, we find, in S/FT networks, during the Cue Off condition, network activity naturally settled onto a stable cyclic trajectory (limit cycle). Limit cycles, however, were a prominent feature only in networks that had undergone the shaping procedure (Only Shaping and S/FT). When the cue switched on, network dynamics shifted toward a strong fixed point that pulled the activity away from the limit cycle (Figure 4a). Switching the cue on or off toggled the active attractor structure, fully accounting for the task-dependent changes in network state. This attractor switching provided a simple and reliable mechanism for flexibly controlling behavior based on task cues.

**Figure 4.**
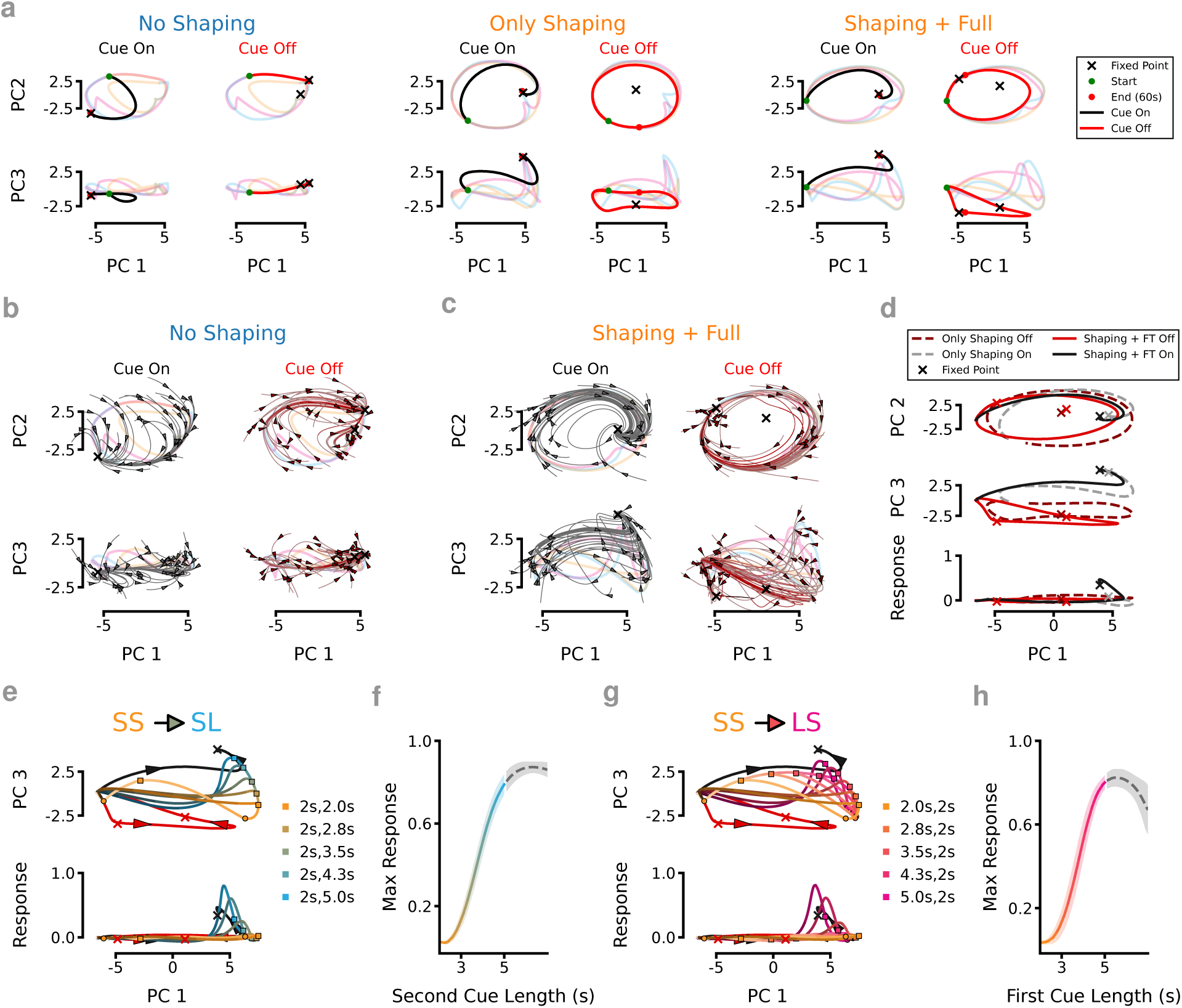
During shaping, networks learn distinct dynamical mechanisms that improve task performance. (a) Evolution of the network state in PCA space for Cue On and Cue Off conditions superimposed over the activity during the full task. Without shaping (*left*), the network dynamics are captured by two opposing trajectories depending on whether the timed cue is on or off. Following shaping tasks (*middle*), a limit cycle is observed which is further refined following training on the full task (*right*). (b) Example PCA state-space flow field for Cue On (*left*) and Cue Off (*right*) conditions for a network which did not get pre-trained on the shaping task. (c) Same as b except for an RNN which was exposed to the shaping task prior to training on the full task. (d) Comparison of the dynamical states learned by an RNN following training on the shaping task (*dashed lines*) and following training on the full task (*solid lines*). Subspaces spanned by PC 1 and PC 2 (*top*), PC 3 (*middle*) and the Response/output node (*bottom*) are shown. (e) Extending the length of the second cue from 2 to 5 seconds forces the network into the response potent space once the duration is sufficiently longer than the ”short” 2s cue length (between 2.8s and 3.5s). (f) Mean and standard deviation of *n* = 13 S/FT RNNs max response value for different cue lengths between SS and SL trials as well as longer-than-long trials (*gray*). (g) Same as e except the first timed cue is varied between the S (2s) and L (5s) duration. (h) Same as f except the first cue length is tested.

In NS networks, we observed two regions contain attractor points that occupy opposite extremes along the main axis of variance (PC1), corresponding to Cue On and Cue Off states (Figure 4a,b, S3a). This geometry pulls network activity along one of two approximate line attractors, depending on the cue state. During Cue On, activity drifts toward a fixed point positioned within the output-potent space, meaning the longer a cue is presented, the greater the likelihood of triggering a response. During the ISI, and inter-trial periods, activity recedes away from the output-potent space. Thus, NS networks behaved like leaky evidence accumulators, where prolonged cues gradually increased response probability. However, network stability in NS networks was poor: input noise during LS trials often pushed activity prematurely toward the Cue On fixed point, leading to aberrant early responses (Figure S4a). We validated this by showing a direct relationship between proximity to the fixed point at the offset of the first cue and the amplitude of erroneous responses (Figure S4b,c). Because the response magnitude of these errors overlapped with correct Go responses, it is impossible to define a clear decision boundary. In contrast, while S/FT networks also approached the Cue On fixed point, they suppressed premature responses due to additional dynamical constraints learned during shaping (Figure S4d-f).

After shaping (and even before full task training), S/FT networks developed critical structure: a stable limit cycle during Cue Off epochs and a fixed point during Cue On epochs offset from that limit cycle (Figures 4a, S3b). This organization already separated response dynamics from basic timing information, preventing simple drift-to-threshold behavior. Following full task training, these dynamics were further refined: the Cue Off limit cycle flattened, keeping network activity away from response-potent dimensions unless specific sequences of Cue On and Cue Off periods had been completed (Figures 4a,c, S3c). To test the generalization of these motifs, we systematically modified cue durations and examined RNN responses. In S/FT networks, activity predictably alternated between the Cue Off limit cycle and the Cue On fixed point, with the network’s proximity to the fixed point—especially along PC3—scaling the strength of the output (Figure 4e). This structured mapping between internal network state and behavioral output held robustly across novel cue manipulations (n = 13 RNNs; Figure 4f). Further, varying the duration of the first cue revealed that the initial cue exposure shifted the starting state for the second cue, effectively gating whether the network entered a response-potent trajectory (Figure 4g,h). Critically, the correct sequencing of Cue On and Cue Off epochs—not just the long cue duration alone—was necessary for generating responses: if the first cue was omitted, network activity remained on the Cue Off limit cycle and no response occurred (Figure S7a). Thus, the networks learned not just isolated stimulus-response mappings, but structured trajectories through state space that dynamically linked prior experience to future behavior.

### Choice of Shaping Task Critical for Learning Abstraction

Learning is often proposed to be compositional (Chomsky 2017, Lee et al. 2024); and previous training episodes are thought to make up building blocks that can be assembled in the future to construct strategies for solving more complex tasks. In fact, it has been shown that training RNNs on very simple tasks can alter the solutions during future learning by constructing important landmarks in the dynamical landscape (Driscoll et al. 2024, Turner & Barak 2023). Therefore we wondered: *does the choice of shaping task matter?* Specifically, if we train an RNN on any of subset of the task, do we alter the network in ways that may improve future learning on the full task? To investigate this, we trained RNNs on a modified shaping task that only included one of the no-match trial types to see if that would similarly aid learning. When we train RNNs on the SL trial type only during shaping (SL/FT networks), we find no significant improvement over networks trained without any shaping phase (Figure 5a,b,h). The SL/FT RNNs adopt the predictable strategy of responding following any long cue (Figure 5c) which leaves them with a sub-optimal solution during the full task that is also hard to refine with additional training. Interestingly, we observe that these SL/FT RNNs follow similar activity patterns during the full task as the NS ones (Figure 5d, S5).

**Figure 5.**
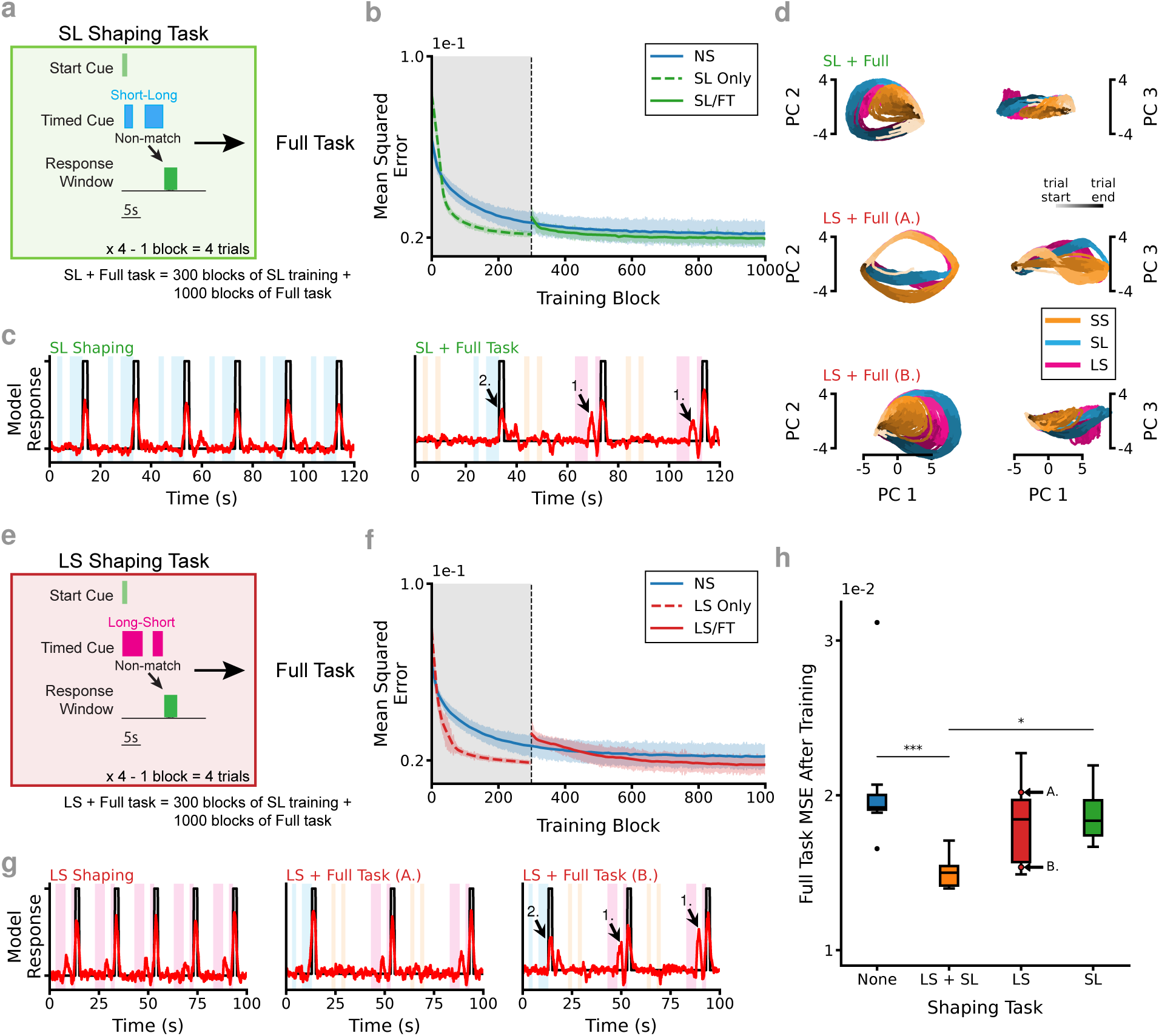
Choice of shaping task dictates networks’ ability to learn necessary abstraction. (**a**) Schematic of shaping on SL trials only (SL/FT RNNs). (**b**) Model performance plateaus after training at similar level as not including the shaping task. (**c**) *Left*. Example of the model response after 300 blocks of the SL Only task. *Right*. Example response after 700 blocks of training on the full task. Note the early responses on LS trials (1.) and the relatively low amplitude response on SL (2.). (**d**) Representation of network activity following training including different shaping tasks. *Top*. Training on SL only results in similar population activity as not including any shaping. *Middle/Bottom*. Training on LS only results in variable outcomes with some networks (A.) able to perform the task while others (B.) are not able to perform the task. (**e**) Schematic of shaping on LS trials only (LS/FT RNNs). (**f**) Model performance plateaus after training but is variable across models. (**g**) *Left*. Example of the model responses after 300 blocks of the shaping task. *Middle* High performing model following LS Only shaping and 700 blocks of training on the full task. *Right* Example low performing model after LS Only shaping and full task training. Note the RNN that makes errors shows early responses on LS trials (1.) and the relatively low amplitude response on SL (2.). (**h**) MSE on full task after training is complete. *n* = 12 NS, 12 S/FT, 11 LS only, 10 SL only, Tukey’s HSD *p−*values as indicated). *-*p <* 0.05, **-*p <* 0.01, ***-*p <* 0.001.

Alternatively, we also train the RNNs on the LS trial type only during shaping (LS/FT networks), prior to training on the full task (Figure 5e). In this case we get a roughly bimodal response where some RNNs perform the task almost as well as the RNNs with the standard no-match shaping procedure while many fail to find a solution and have performance similar to SL/FT and NS networks (Figure 5f,g). We observe remarkably different structure in the PCA state space plots between RNNs with these two results. The high performing LS/FT networks form a ring structure which is conserved between the three trial types, while the LS/FT networks that do not perform the task well do not (Figure 5d). This means that based on the randomized initialization of the networks and the noise present during training, some networks learned to generalize the temporal structure between trial types and thus were able to perform the task well, while others did not learn the abstraction. This validates our central hypothesis that the shaping task embeds critical structure into the networks that aid future learning, but, *why does this happen sometimes for LS/FT networks and never on the SL/FT ones?* Critically, the added complexity of having to identify the longer cue (which indicates that this is a Go trial), but then waiting until after the second cue to respond, requires a more nuanced strategy and may sometimes be sufficient to train RNNs to perform the full task well. More generally, as it relates to the tDNMS task, we observe that shaping on only the SL or LS trial type is sub-optimal compared to shaping on both of the no-match types as the RNNs fail to robustly learn an optimal solution to the tDNMS task (Figure 5h). The inclusion of two trial types exposes the networks to multiple temporal contexts during the pre-training phase of learning, forcing them to learn the abstraction that is necessary to adopt the more optimal strategy.

### RNN predictions demonstrate how shaping procedures are critical to the development of key task abstractions in animals

The tDNMS task was originally designed as an odor timing task for mice (Bigus et al. 2024). Analysis of the RNNs is most interesting if it gives insight into the behavioral strategies and neural dynamics that animals develop as they learn and perform the task. To test the predictions of our RNN-based analysis in animals, we trained a cohort of mice on the SL-Only shaping task rather than the standard no-match procedure. We then tracked mouse behavior over 8 days, a time sufficient for most animals to reach proficiency in the task (Bigus et al. 2024). We selected the SL-Only shaping since the RNNs make specific testable predictions about how the strategy would be altered in this case. The RNN analysis predicts that the SL-Only shaping procedure should develop strategies on the full task that closely match the NS RNNs (Figure S5), while providing animals with a task that familiarizes them with the recording setup, reward port and cue-ISI-cue-respond task structure. After the animals reached proficiency in the shaping task (*see Methods*), we performed surgery to chronically implant Neuropixels 2.0 probes (Steinmetz et al. 2021, Melin et al. 2023) and then tracked the behavior and neural activity of the animals on the full task (Figure 6a, S6a,b).

**Figure 6.**
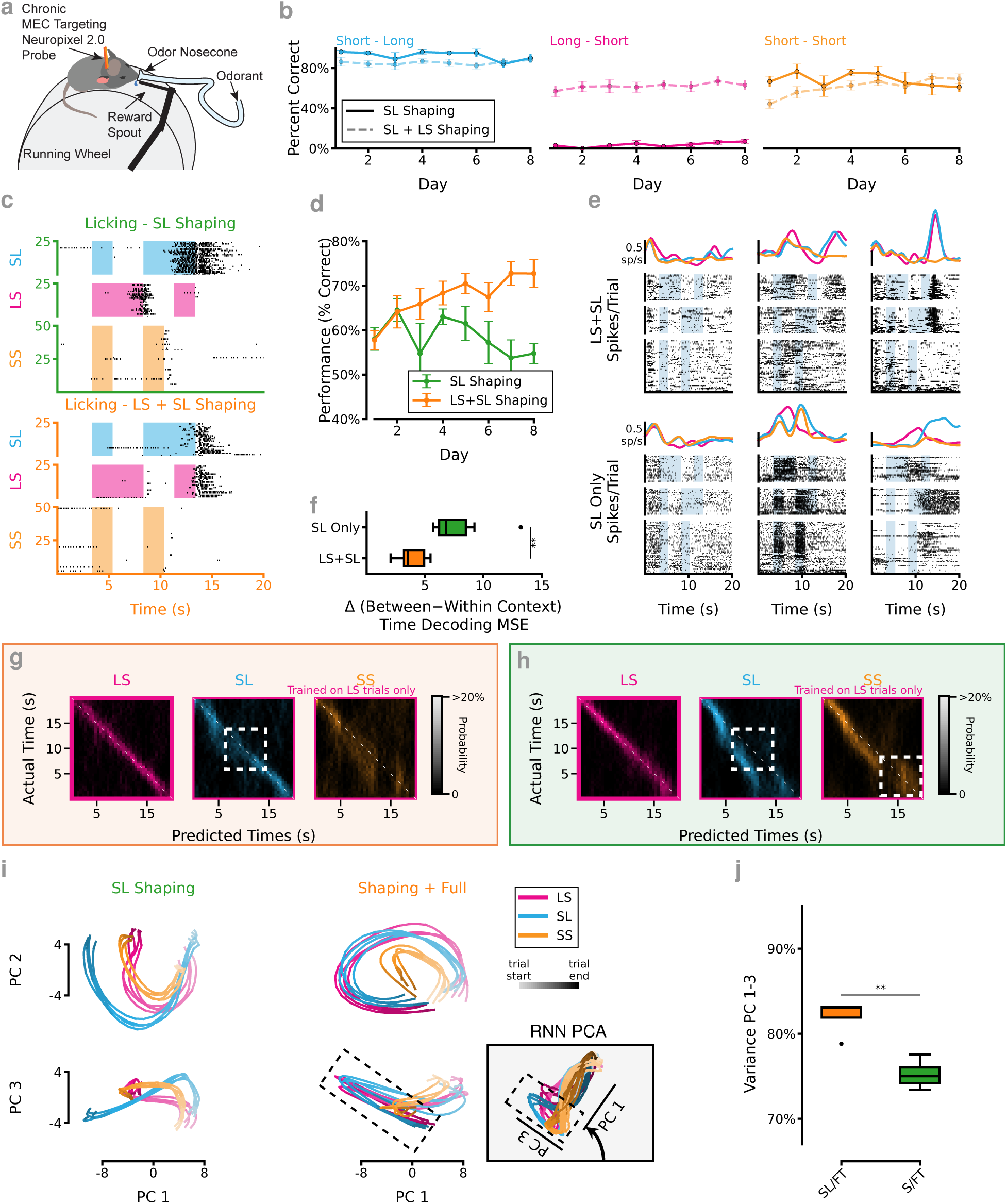
Past experience predicts abstraction learning in animals performing tDNMS task. (**a**) Mice perform an analogous cue timing task with an odor cue and and licking for water rewards. Separate cohorts were trained with SL + LS Shaping and SL Only shaping. (**b**) Comparison of animal performance by trial type based on shaping experience across the first 8 days of Full Task training. *n* = 16 mice SL+LS shaping, *n* = 4 mice SL Only shaping. (**c**) Example licking behavior of a well trained mouse on the Full Task. *T* op. After SL Only shaping. *B*ottom. After standard LS+SL shaping. (**d**) Performance averaged across all trial types. *n* = 16 mice SL+LS shaping, *n* = 4 mice SL Only shaping. SL Only mice are significantly worse after 8 days of training *p* = 0.002 (**e**) Example spike raster and tuning curves for mice recorded during the Full Task. *T* op. For an animal after LS+SL shaping. *B*ottom. For an animal that went through SL Only shaping. (**f**) Error comparison decoding trial time from animal MEC based on shaping experience. Similar to the RNN trained without proper shaping, SL Only trained mice show decreased performance at cross-context time decoding, *p* = 8.72 *×* 10*^−^*^3^, *n* = 8 SL Only mice, 4 mice x 2 session / mouse, *n* = 5 sessions, 4 mice SL+LS Shaping. (**g**) Confusion matrix for time decoding on mice trained with LS+SL shaping (10-fold cross validated) decoder trained on LS trials then tested on LS (left), SL (middle), and SS trials (right). Average of *n* = 5 sessions from 4 mice. (**h**) Same as g for SL Only trained animals. Average of *n* = 6 sessions from 3 mice. (**i**) PCA projection of pseudo-population task aligned units recorded from MEC during tDNMS task. *Left*. Animals traing on SL Only shaping prior to introducing the full task. *Right*. Animals with SL+LS shaping (S/FT). *Top*. Plot of PC1 and PC2. *Bottom*. Plot of PC1 and PC3. *Inset*. S/FT RNN prediction for comparison. Dashed lines indicate response window. (**j**) PCA projection of the pseudo-population task aligned units shows increase percentage of variance in the first 3 PCs from animals with SL+LS shaping. *n* = 4 random splits of pseudo-population data, *p* = 2.73 *×* 10*^−^*^3^ two sample independent t-test.

We first sought to answer the question: *Do animals also learn the same abstracted temporal representations as observed in the RNNs?* To answer this, we first compared the behavior of animals trained on the SL-Only shaping to animals trained with the standard no-match shaping task. We find that the SL-Only shaping animals are unable to learn to adapt their behavior to find an optimal solution to the task, perseverating on a “lick-after-long” strategy (Figure S6c,d). Which, while not a solution capable of gaining the maximum possible rewards on the task, is still capable of getting 75% of the trials correct–all of the SL and SS trials and none of the LS ones (Figure 6b,c,d). This closely matches the response of both the SL/FT RNNs (Figure 5c) and the most common errors made by NS networks (Figure 1h,S4).

The RNN model predicts that proper shaping tasks should embed critical between-trial type abstractions about the structure of the task. We validated this in the RNNs by showing improved between-context decoding in RNNs trained on the proper shaping task while NS and SL/FT networks have much worse performance on this task (Figure 2d,S5). We target Neuropixels recording to the MEC since it was previously shown that the MEC is necessary to learn successful strategies for performing the tDNMS task and choice of strategy is reflected in MEC activity (Bigus et al. 2024). Additionally, MEC is a highly recurrently connected brain region with computational mechanisms that could have strong parallels to the RNN model (Zutshi et al. 2018). In line with previous results, we find cells in the MEC that appear to have trial-time locked tuned fields (Figure 6e). Performing both within- and between-context trial time decoding, we find that the mice trained on the LS-Only shaping procedure have much worse between-context decoding compared to within-context while mice with the standard no-shaping procedure have more similar decoding errors in both cases (*p* = 8.72 × 10^−3^, two-sample t-test, Figure 6f). In fact, when we look at the errors made by the time decoding we see very similar error patterns on the cross-context decoding test as predicted by the RNNs (Figure 2d, 6g,h, S6f,g).

In order to visualize how the population activity of within MEC during the task, we generated a pseudo-population of task-responding neurons (*n* = 384 units, Pearson’s *ρ >* 0.25) and calculated the first three PCs (see Methods). We find remarkably different responses between mice trained in SL-only shaping animals (SL / FT)) compared to animals trained in S / FT FT compared to standard shaping (S/FT, Figure 6i). Additionally, the S/FT trained animals show structured dynamics in PCA space similar to the S/FT network, subject to rotations of the state space, with circular representation in the first two PCs and response valence visible in the plot of PC1 and PC3. Lastly, we look at the percentage of explained variance in the first three principal components and find that the S/FT mice have a similarly lower dimensional representation of the state space compared to SL/FT trained animals (*p* = 2.73 × 10^−3^, *n* = 4 splits of pseudo-population data for both SL/FT and S/FT trained animals, Figure 6j) similar to the RNN model (Figure 2b). We also find instances of higher tangling in MEC dynamics during the tDNMS task in SL/FT trained mice than S/FT (Figure S6e). These results demonstrate that using RNNs to model the underlying dynamics that evolve with experience has permitted us not only to resolve predictions about the behavior of animals, but also to discover essential aspects of the neural computations supporting task performance. Particularly, we make these observations in a brain region that has previously been shown to be critical to strategy development.

### RNNs correctly predict behavioral responses to novel trial types

A critical observation of the dynamical mechanisms responsible for the strategy employed by S/FT networks is that the Cue On fixed point is in the output-potent space of the RNN. Further, peak responses happen as the activity falls from the fixed point back to the Cue Off limit cycle (Figure 4). This means that, while the timing of the cues is necessary to get the RNN to respond, there’s a larger set of cue-timing configurations that would result in a response beyond the two “Go” trial types that the networks were trained on. Specifically, the RNN predicts that both a Long-Long (LL) and a Medium-Medium (MM) trial would result in a response (Figure 7c, S7b). Interestingly, this is at odds with the explicit Non-Match ≡ “Go” paradigm that is implied by the Non-Match to Sample task. As such, we validated this behavior with animals well-trained on the tDNMS task. In order to test animals on the probe (LL or MM) trials without encouraging them to adjust strategy, we implemented a modified tDNMS task where 20% of trials were unrewarded, novel probes (Figure 7a). Importantly, animals maintain proficiency at the tDNMS trials, and with 80% of the trials being the familiar trial types, mice continue to lick robustly in the response windows and withhold licking on the “No-Go”/SS trials (Figure 7b,d). Additionally, animals do indeed lick in response to both the LL and MM trials (Figure 7b,e,f). Previous modeling of the animals’ behavior indicates that mice use the presence of the Long cue as an indication that they are in a “Go” context while timing both cues to know when to respond (Bigus et al. 2024). As such, the LL response is expected–and MM response is only dependent on how much longer-than-short the cue is before animals interpret the cue as being long (e.g. how optimistic the animals are). The RNN analysis predicts that any cue longer than around 2.8s would be long enough to elicit a response (Figure 4e-h).

**Figure 7.**
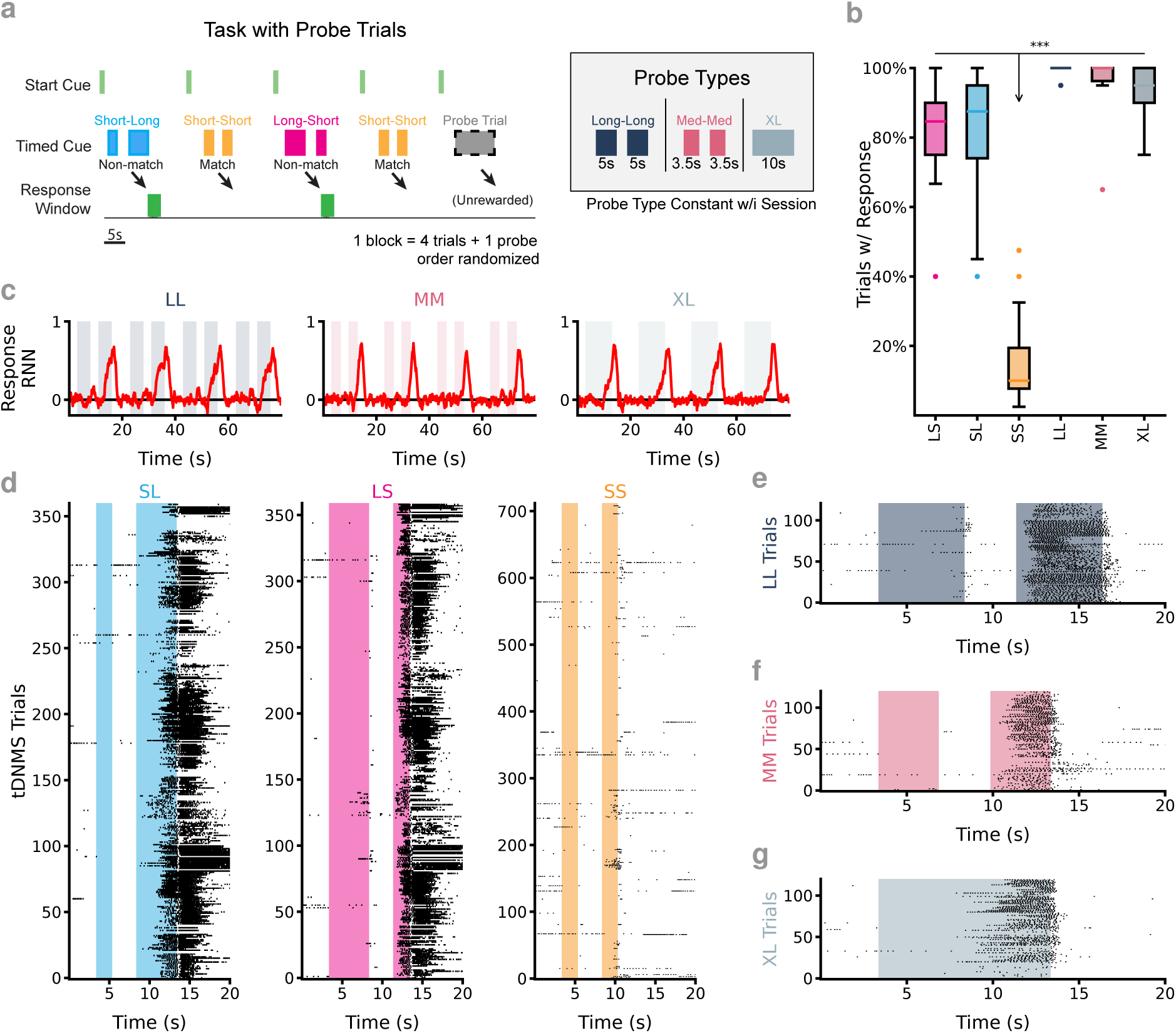
RNNs predict animal responses to novel temporal contexts. (**a**) In order to test behavioral response to previously untrained trial types (temporal context), novel cue durations were interleaved into the block structure of the tDNMS task. On each day, a single probe trial was selected and presented to the animal on 20% of trials. Probe trials were unrewarded so as not to reinforce the decision made by the animals. (**b**) Animals robustly interpret probe trials as a “Go” trial type, responding significantly more than to the SS/“No-Go” trial type. *n* = 6 probe days for each probe type from *n* = 3 mice and *n* = 18 non-probe sessions (pooled across all probe recording days). 2 Factor ANOVA, significant effect of trial type, *p* = 1.03 *×* 10*^−^*^22^, mouse, *p* = 0.02, but not the interaction between mouse and trial type *p* = 0.50. Response probabilities significantly grater than “No-Go” trials: LL,SS *p* = 1.24 *×* 10*^−^*^14^, LS,SS *p* = 1.24 *×* 10*^−^*^14^, MM,SS *p* = 1.27 *×* 10*^−^*^14^, SL,SS *p* = 1.24 *×* 10*^−^*^14^, XL,SS *p* = 1.29 *×* 10*^−^*^14^. All others not significant, Tukey’s honestly significant difference test for multiple comparisons. (**c**) After training, S/FT RNNs robustly respond following the offset of the second cue in each of the three probe trial types. (**d**) All licking behavior during probe sessions. Mice continue to lick in the response window during tDNMS non-match trials when unchanged probe trials are interleaved. Data from 3 mice, each recorded 6 times (2 per probe). (**e**) Mice lick in response the LL probes. Data from 3 mice recorded in 2 separate sessions per mouse. (**f**) Mice lick in response the MM probes. Data from 3 mice recorded in 2 separate session per mouse. (**g**) Mice lick in response the XL probes. Data from 3 mice recorded in 2 separate session per mouse.

Interestingly, there is only a single fixed point in the Cue On mode that the networks are pulled toward. This, surprisingly, implies that a sufficiently long single cue should result in a response. We validate this prediction by testing it with S/FT RNNs and find that a single, 10s, extra-long (XL) cue results in a response and find that RNNs reliably respond to this trial type (Figure 7c). We also validate this behavior in animals through a third probe trial session and find that they do indeed lick in response to XL trials (Figure 7b,g). We consider this a somewhat surprising result as animals (and RNNs) are always trained with the two-cue trial structure and cue transitions are likely more salient than timing a persistent cue. The remarkable fidelity of animals to the strategy predicted by the RNNs further validates the evident parity between the mechanisms governing the RNN dynamics and how the animals are solving the tDNMS task.

## Discussion

Using a complex odor timing task, we show that both RNNs and animals benefit from having a carefully structured prior learning experience. In particular, training with a no-match shaping paradigm enables the system to learn key abstractions that improve later performance on a more complex task. We specifically designed the tDNMS task to require cognitive demands beyond simple stimulus-response learning, allowing us to examine how early experience influences the strategies animals and networks construct. In motor learning, the building blocks–such as the steps of a dance–are often clear; in contrast, in cognitive tasks like the tDNMS task, animals must learn to integrate information about the cue durations and sequence over extended time periods and across gaps (the ISI) to infer whether they are in a Go or No-Go context. This task complexity allowed us to isolate how prior shaping influences learning in environments where context must be inferred without an explicit cue. In our task we have 3 trial types that portend 2 possible outcomes, all based on a single input stream. Animals must recognize the current trial type through real-time observation of the cue and store that information to guide future actions–a process analogous to everyday situations where appropriate behavior depends on inferred, overlapping contexts. We show that prior experience constrains the state space that a dynamical system traverses to be low dimensional, resulting in the learning of key abstractions about the commonalities between the trial types. This abstraction simplifies subsequent decision-making. Together, our results identify a mechanism for curriculum learning in complex cognitive tasks–one that does not depend on explicit contextual cues or fully orthogonal input spaces, distinguishing this work from many previous studies on learning dynamics and multi-task training (Driscoll et al. 2024, Mante et al. 2013, Yang et al. 2019, Lin et al. 2023).

Notably, the strategies adopted by the RNN models share much in common with the strategies adopted by animals performing the task. We see this demonstrated in both the behavioral predictions from altered shaping experience as well as in testing animals with ”probe” trial types to validate performance on previously unseen temporal structures. For example, we had assumed that the presence of both odor cues is an integral part of the trial structure that animals would follow in order to assess proximity to the response window. However, the S/FT RNN strategy does not depend on the ISI or counting independent odors, a notable insight that led us to test, and confirm, in animal behavior. Ultimately, we also see reflection of cognitive strategy in the neural activity recorded in MEC. Mice undergoing the SL-Only shaping procedure have skewed task representations that match the NS or SL/FT RNNs (Figure 2d,6h). The mice trained on the no-match shaping performed the tasks well and show evidence for generalization between trial types (Figure 2f,6g). Thus, prior experience on the (correct) shaping task allows animals to recognize key similarities in the structure of the trial, resulting in more low dimensional task representations. We see the past experience as having built a conceptual scaffold, that constrains future activity to traverse prescribed states. When novel task demands are then placed on the system, the natural solutions are to move in directions orthogonal to that scaffolding, resulting in the observed separation of the time-decoding generalization and response-generating states into orthogonal dimensions. Importantly, these insights were only possible because of our theory-driven approach: using RNN models to generate specific, mechanistic predictions about behavior and neural dynamics, and then directly testing those predictions through large-scale neural recordings in behaving animals. This closed-loop model-experiment pipeline allowed us not only to confirm that the general principles of abstraction learning apply to both artificial and biological systems, but also to uncover dynamical motifs that would have been extremely difficult to detect without targeted hypotheses. Our approach thus provides a framework for understanding how structured experience shapes learning strategies at both behavioral and circuit levels.

Curriculum learning is often posed as a problem of training a system on the correct task primitives that can be assembled, like building blocks, into a strategy for solving a novel, or more complex task. While our results are not strictly in disagreement with this idea, we demonstrate that for complex tasks - specifically those requiring context-dependent cognitive strategies - the correct task building blocks must be identified carefully. Training RNNs or animals on SL-Only shaping results in a suboptimal strategy that is hard to unlearn, however, SL trials are explicitly a sub-task of the tDNMS full task. This proposes an exciting avenue for future experimental research. As we desire to train animals on increasingly complex and ethologically relevant tasks (in which animals build up a repertoire of relevant experience over their adolescence), it becomes harder to ad-hoc design shaping curriculum that support the task structure or guess at what strategy they will arrive at to perform the task. Resolving the strategies adopted by a simulated dynamical system gives important clues as to how a recurrently connected dynamical system (such as an RNN or the brain) might build up the necessary lower dimensional structures leading to appropriate task chunking in a way that novel task elements are easier to learn. Understanding how early training experiences shape these internal structures could also have important implications for optimizing learning in aging populations or in neurodegenerative diseases, where building effective strategies from limited plasticity and degraded representations may be especially critical. Previous results (Hocker et al. 2024) as well as ours (Figure 6c,f,g,h) indicate that appropriate pre-training steps cause artificial learning systems to adopt strategies that more closely match with those that animals adopt. The cost in time and resources is dramatically less for training RNNs then performing animal research. Since developing training strategies in simple models and fitting to existing observations allows us to reason about how future tasks will be learned, it is possible to test out curriculum *in silico* before moving to *in vivo* experiments.

It is important to consider that we did not fit biophysical models to neuronal data or employ ethologically plausible learning rules. Instead, we designed a simple model capable of performing our task, subject to constraints on noise and regularization terms, and used it to make predictions about how animals would solve the task. We then validated specific predictions about neural dynamics and behavior. The parameterization of the RNNs is held consistent in order to get fair comparisons between models based on experience. Another important consideration is that the RNNs we train perform only a single task and we do not consider the case of “catastrophic” forgetting (Duncker et al. 2020, Kirkpatrick et al. 2017). Since our objective is to understand how performance on a single task evolves with experience, we do not consider the case of multi-task learning. Particularly, in our investigations, the shaping tasks we test are all subsets of the full tDNMS task. Expanding this work to how multiple distinct tasks overlap in their representations and alter the strategies adopted is another key area for future research, especially in the case when contextual information is necessary to perform the correct actions and context needs to be inferred from the cues in the environment. Finally, we show that past experiences can both amplify performance as well as interfere with finding optimal strategies. Investigating how these interactions reach across task domains could suggest potential mechanism for improved learning strategies or offline learning that could be transferred to support more optimal performance on future complex tasks.

## Acknowledgments

We thank Brian DePasquale and Laura Driscoll for their valuable comments and feedback on earlier versions of this manuscript. The present study was supported by the U.S. National Institutes of Health (NIH), the National Science Foundation (NSF), the NIH/National Institute of Mental Health (grant no. 1 DP2 MH129958-01 to J.G.H.), an NSF CAREER Award (no. IOS-2145814 to J.G.H.), the Life Sciences Research Foundation with generous support from the Simons Foundation – Collaboration on the Global Brain (J.C.B.), and the University of Utah (J.G.H.). The funders had no role in study design, data collection and analysis, decision to publish, or preparation of the manuscript.

## Author Contributions

J.C.B. and J.G.H. designed experiments and analysis, and wrote the manuscript. J.C.B and D.A. developed and the RNN model and analysis. J.C.B., D.A., C.J. and H.L. collected, analyzed and interpreted the data from animal experiments.

## Methods

### Lead Contact and Materials Availability

Further information and requests for resources and reagents should be directed to the Lead Contact James Heys.

### Experimental Model and Subject Details

All experiments were conducted in accordance with the NIH guidelines and with the approval of the University of Utah Institutional Animal Care and Use Committee. Experiments were performed with adult (8-16 weeks) C57Bl/6 mice (Charles River Laboratory).

## Methods Detail

### RNN implementation and training

To train RNNs to perform the tDNMS task, we used a modified implementation of a “SimpleRNN” from the python package tensorflow (Abadi et al. 2015) that was edited to provide for a “leak” term to force the activity of the individual units to decay over time (in the absence of any input) as well as to implement a private noise for all units in the RNN. The activity of the networks are therefore governed by the equation:

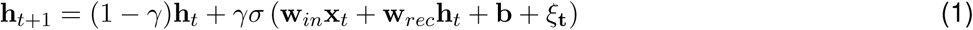

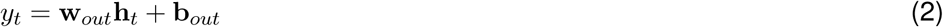

Wherein *γ* is the added activity decay bias, **x***_t_* is the 2 × *T* input values, **w***_in_* is the 2 × *N* input values, **b** unit biases, *ξ_t_* private unit noise, **h***_t_* unit activations on the current time step, and **w***_rec_ N* × *N* connection weights in the recurrent layer. *σ* represents the non-linear activation function which was tanh for all RNNs tested. To determine the value of the response node, **y***_t_*, **w***_out_ N* × 1 output weights, and, **b***_out_*, the output bias were used. For all RNNs *N* = 128 units, the constant *γ* = 0.2 and *ξ_t_* was drawn randomly from N (0, 0.3) on each time step. To prevent overfitting, the inputs were disturbed by adding noise randomly draw from N (0, 0.15). **w_in_**, **w_rec_ b**, **w_out_**and **b_out_** were learned while training the RNNs.

To train all RNNs, we used tensorflow’s implementation of backpropagation through time using the proved implementation of the Adam optimizer to preform adaptive gradient descent when determining weight updates (Kingma & Ba 2017). For training, *L*_2_ regularization was applied to both the weights and activations with *λ* = 1 × 10^−3^ and Adam learning rate set as *α* = 1 × 10^−5^ and a limited batch size of 2 was used. RNNs were initialized such that the input weights were drawn from a uniform distribution and the recurrent layer was initialized to be a random orthogonal matrix. Each RNN was initialized separately and trained with unique input and unit noise at each time step. Additionally, the order of trial types was randomly shuffled within blocks of 4 trials, independently every time the RNN was run. All RNNs were trained for a total of 1000 blocks of 4 trials, if shaping was used it replaced the first 300-4 trial blocks. For training and analysis, time was discretized into 10 ms bins, such that there were 200 time bins per 20s trial.

### RNN Analysis

#### Time Decoding

Time decoding was conducted using a Linear Discriminant Analysis (LDA) based classifier implemented in scikit-learn (Pedregosa et al. 2011). We first generated 60 trials worth of data (15 × 4 blocks of trials), then split the resulting activity of the recurrent layer of the RNN into 10 folds for cross validation (by trial). To calculate within- vs across-context decoding scores, we trained 4 decoders for each split–one including only trials from 1 of the 3 trial types in the training set and a fourth with all trial types. We then tested the decoders on the held out data from all trials and report within-context decoding as the subset of trials when the training decoder matched the testing trial type and across-context decoding otherwise and predicted elapsed trial time to the nearest 1s. The trained on all trials decoder was only used in Figure 2e for comparison.

#### Trajectory Tangling Metric

Tangling is a metric to quantify the degree of smoothness a trajectory takes in state space and it has been shown that avoiding tangling is an important property of neurons in the motor cortex (Russo et al. 2018). High tangling indicates that similar locations in state space are re-purposed to represent distinct moments within or between temporal contexts. The tangling at each time step, *Q* (*t*) is calculated as:

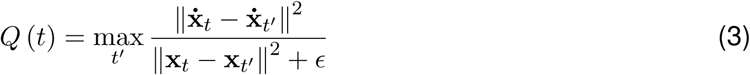

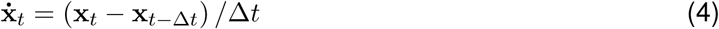

Where **x_t_**is the state at time *t* and **x_t_** is the derivative (calculated numerically as in Equation 4) with Δ*t* = 0.1 and *ɛ* was set to 0.1 times the average squared magnitude of **x_t_**for all values of *t*.

#### Fixed Point and Flow Field Analysis

We identify fixed points according to the methods developed in Sussillo & Barak 2013 and Golub & Sussillo 2018 and most closely resembling the techniques used to identify dynamic motifs in Driscoll et al. 2024. We use a modified version of the fixed point finder code (https://github.com/google-research/computation-thru-dynamics, Apache-2.0 license) implemented in Python using the *JAX* library (Bradbury et al. 2018). Briefly, this method identifies candidate fixed or slow points in the RNNs’ state space based on the trajectories traversed during the task and then uses an Adam based optimizer to perform gradient descent to find where the network settles. This is possible because around the fixed points we can approximate the evolution of non-linear RNN dynamics as a linear system using Taylor series expansion, around candidate fixed points, **h**^∗^:

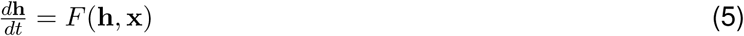

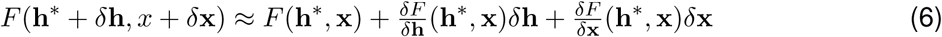

Particularly, we treat Cue On epochs and Cue Off epoch separately, meaning that *δx* = 0 in all cases and since ∥*δh*∥*^n^* ≈ 0 for *n >* 1 the higher order terms in the Taylor series can be ignored.

Therefore, the linearized system (near fixed points) is approximated as *<u>^δF^</u>* ≈ *<u>^δF^</u>* (**h**^∗^, **x**)*δ***h**. When identifying fixed points, candidate points are generated based on the the states occupied by the RNNs during the full tDNMS task and set the input and internal noise terms to 0. Lastly, in addition to finding fixed points, we also identify points that are sufficiently slow on the timescale of the tDNMS task as approximate fixed points or slow points and define squared speed as *q*(**h**) = <u>^1^</u> |*F* (**h**)|^2^, and label points with *q <* 10^−6^ as fixed points for our analysis. We validated the fixed points and identified limit cycles by permitting networks to run for long durations (2× trial length) and find this method sufficient to identify the dynamical motifs governing the strategies adopted by the RNNs in our analysis.

### Animal Surgery and Recordings

#### Neuropixels Implant

A Neuropixels 2.0 probe (NP2014, Imec), encased in a 3D-printed part (Melin et al. 2023), was disinfected with 99% isopropyl alcohol (IPA) for 1–2 minutes. The four shanks of the probe were dipped ten times into a Vybrant cell-labeling solution (Thermo Fisher) to mark the probe track in the brain. C57BL/6 mice (Charles River Laboratory), aged 2–3 months, were chronically implanted with Neuropixels probes. Mice were anesthetized with 1 − 2% isoflurane and secured using ear bars and a tooth bar in a stereotaxic frame (Kopf Instruments). Eyes were protected with ophthalmic ointment. Fur over the scalp was removed using depilatory cream, and the scalp was sterilized with ethanol and iodine. An incision was made to expose the skull. Small holes for skull screws were drilled on the left hemisphere and over the cerebellum. One screw was inserted into the left hemisphere, and another screw connected to a silver wire was inserted into the cerebellum; the silver wire served as a ground for neural recordings. A mark for craniotomy was made by black ink at 3.3 mm lateral to bregma around the lambdoid suture line. A custom-designed titanium headplate (9.5 × 33 mm²) was placed on the dorsal surface of the skull, and an opaque Metabond (Parkell) was applied on the skull to fix the skull screws and headplate. After recovery from the surgery, mice began to train in the tDNMS task.

Once mice were proficient at the shaping task, a second surgery was performed. Anesthesia and fixation procedures were identical to the first surgery. The opaque Metabond was removed by drilling at the black mark on the skull. A craniotomy (2 mm × 1 mm) was performed on the right hemisphere, centered 3.3 mm lateral to bregma and slightly caudal to the lambdoid suture, exposing the anterior portion of the transverse sinus. A durotomy was performed under sterile saline. After removing the skull fragment, a ring-shaped well was built around the craniotomy using Charisma (Heraeus Kulzer). The Neuropixels probe was inserted at an angle of 11–13 degrees anteriorly. The center of the four shanks was targeted at 3.3 mm lateral to bregma and anterior to the sinus. The probe was slowly lowered at 2*um/s* until reaching either a depth of 3.5 mm or until one of the shanks began to bend. Upon reaching the target depth, dura-gel (Cambridge NeuroTech) was applied to the craniotomy to maintain brain health. Kwik-Sil (World Precision Instruments) was then applied around the probe to seal the shanks, followed by application of Metabond (Parkell) over both the skull and Kwik-Sil to fix the probe case. After each surgery, carprofen (OstiFen™) was administered subcutaneously (5 mg/kg) for pain and inflammation management for two days or until the mouse had fully recovered.

For one mouse recorded under an acute setting rather than chronic implantation, the craniotomy was performed under anesthesia the day before recording. The probe was inserted into the brain while the mouse was awake and head-fixed in the behavioral rig. After each recording session, the probe was retracted, and the same craniotomy was reused for two additional days. Different fluorescent dyes (Vybrant™ Multicolor Cell-Labeling Solutions: DiO, DiI, DiD, Thermo Fisher) were applied to the probe to differentiate recording tracks across days. Between sessions, the craniotomy was covered with Kwik-Sil.

#### Electrophysiology Recording

Neural recordings were processed on a headstage (HS-2010, Imec) where signals were amplified, multiplexed, filtered, and digitized. These signals were then transferred to an acquisition system (Imec and National Instruments) and streamed into SpikeGLX software (https://billkarsh.github.io/SpikeGLX) for recording. Data were sampled at 30 kHz, and digitized with a gain of 100. Behavioral signals, collected using BNC-2110 and PXIe-6341 hardware (National Instruments), were simultaneously streamed into SpikeGLX and time-aligned with the neural data.

#### Single-unit Spike Sorting and Analysis

In order to perform spike sorting and identify putative single units for subsequent analysis, we used the graph-based clustering algorithm implemented in Kilosort 4 (Pachitariu et al. 2024). The Python package spikeinterface was used to pre-process the datasets, interface with Kilosort 4, and apply quality control metrics to the output (Buccino et al. 2020). Recording channels were grouped by shank on the 4-shank neuropixels 2.0 probe and processed independently. We then high pass filtered all channels at 400 Hz, applied phase shift correction and common reference averaging across all channels. Bad channels were identified and removed based on noise estimation and similarity to nearby channels. Motion artifacts were detected and removed using the modular motion detection package built into spikeinterface (Garcia et al. 2024) and the Kilosort-like method (Pachitariu et al. 2024). Lastly, all units included in analysis were required to have a minimum ratio of 0.9, inter-spike-interval violations ratio of less than 0.1 and amplitude cutoff of 0.1 (Hill et al. 2011), and verified that these metrics using the visualization tool Phy2 (https://github.com/cortex-lab/phy).

Time decoding and PCA was performed on the identified single units identically as it was performed on the RNNs (described above). To convert spiking into an analogous rate code to the RNNs, the spike times for each unit were grouped into 0.5s bins and Gaussian smoothing was applied with a *σ* = 2s and a window size of 8s and then z-scored. To generate a pseudo-population for PCA plots, task-responsive units were identified by calculating the mean trial-to-trial Pearson’s correlation. Highly correlated units (*p >* 0.25) were retained the S/FT and SL/FT datasets were down sampled to ensure equal number of units in each. In order to pool data between mice while estimating the variability in the data, representative LS, SL, and SS tuning curves were generated on 4 trial splits of the data and PCA was calculated on the result. The variance explained by the first three principal components was calcuated sperately for each trial split of the resulting pseudo-population.

### Animals Training and Behavior

Previously established protocols for tDNMS training were followed for the experimental portion of this study (Bigus et al. 2024). In summary, training started with habituation, where head-fixed mice were provided with water reward, and after reliably licking *>* 80% of the drops in a series of 50 drops, they began training. Training structure remained the same: 3s before odor onset, a flash of green light cued trial start, then the first odor came on, followed by a three second interstimulus interval, and then the second odor, with a response window following second odor offset. Mice were trained on one session of 100 trials per day for six days a week. Training began with shaping, during which mice in the full shaping protocol were presented with randomly intermixed SL and LS trials, whereas mice in the SL shaping protocol were presented with only SL trials. In shaping phase 1, a drop (5ul) of water was administered after second odor offset, until mice licked to take water in *>* 80% of trials. Progressing to shaping phase 2, mice had to lick in a 3s response window following second odor offset to earn reward. If mice failed to earn reward, the next trial was automatically rewarded. Shaping phase 2 continued until mice successfully triggered reward in *>*20 consecutive trials. Advancing to shaping phase 3, mice had to earn reward by licking during the response window and withhold licking during the first odor presentation and inter-stimulus interval. An incorrect trial was followed by a trial with automatic reward delivery as in shaping phase 2. Behavior was monitored until mice earned *>* 20 consecutive rewards for two consecutive days. Subsequently, mice advanced to the tDNMS task, where all trial types were introduced. Mice were rewarded for non-match trials if they licked in the response window and withheld licking before the second odor onset and received no reward for match trials. For full shaping mice, Neuropixel 2.0 probes were implanted once the animals performed *>* 70% correctness, and neural data was recorded after the recovery. SL shaping mice underwent Neuropixel 2.0 probe implant surgery after the completion of shaping 3. After recovery, mice were trained in the full tDNMS task, which continued until day 8 of the task. Electrophysiology data was recorded on the first three days of the task and then alternating days until training was complete.

#### Probe Trials

A separate cohort of mice that were proficient in the full tDNMS task were trained on a version of the task with varying probe trials. Following a day after performing well on tDNMS, mice underwent a session with blocks of 5 trials: 2 nonmatch, 2 match, and 1 probe trial that were randomly shuffled for each block. Sessions with probe trials consisted of either 3.5s medium odor, 3s ISI, and 3.5 medium odor (MM); 5s long odor, 3s ISI, and 5s long odor (LL); or one extra-long odor of 10s (XL).

After probe trials, mice were returned to the tDNMS task schedule until they performed proficiently again, which would lead to another session of a different probe trial type the following day. All mice underwent each probe trial type twice.

## Supplementary Materials

**Figure S1.**
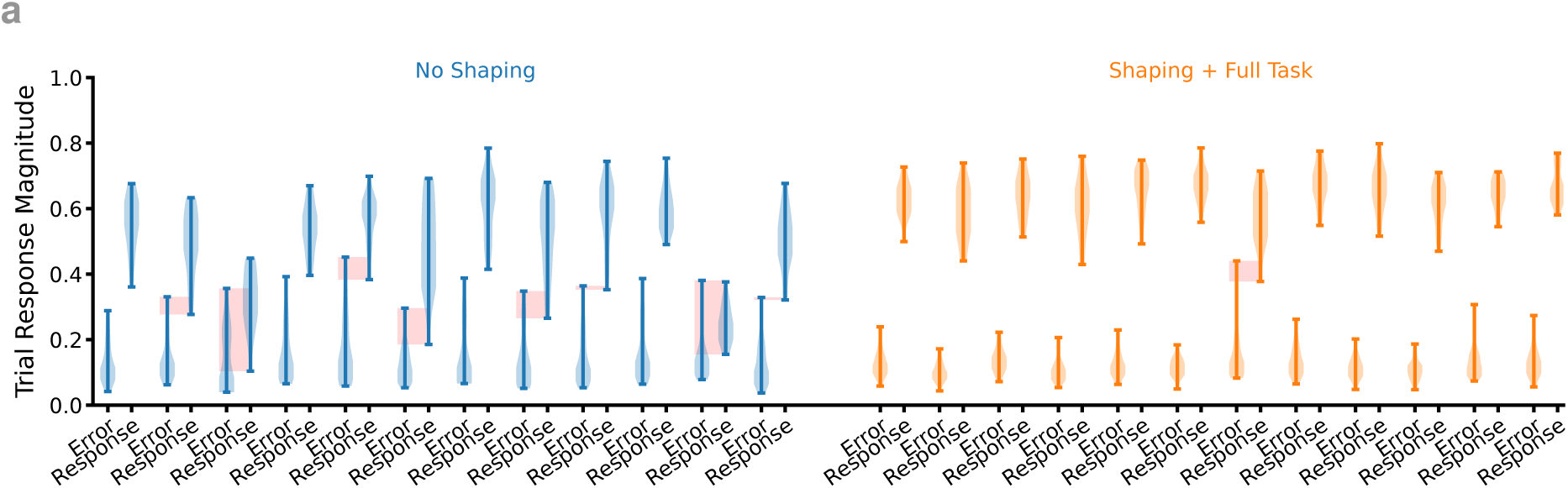
S/FT RNNs more robustly solve the tDNMS task compared to NS RNNs. (**a**) Peak values inside the “Error” window compared to the “Response” window during tDNMS task for *n* = 12 NS and *n* = 12 S/FT networks. Following shaping, individual RNNs solve the task robustly for a range of thresholds on the value of the response node, with many RNNs showing an overlap in response magnitudes between the response window and other times throughout the trial (red boxes) or very narrow window of threshold values.

**Figure S2.**
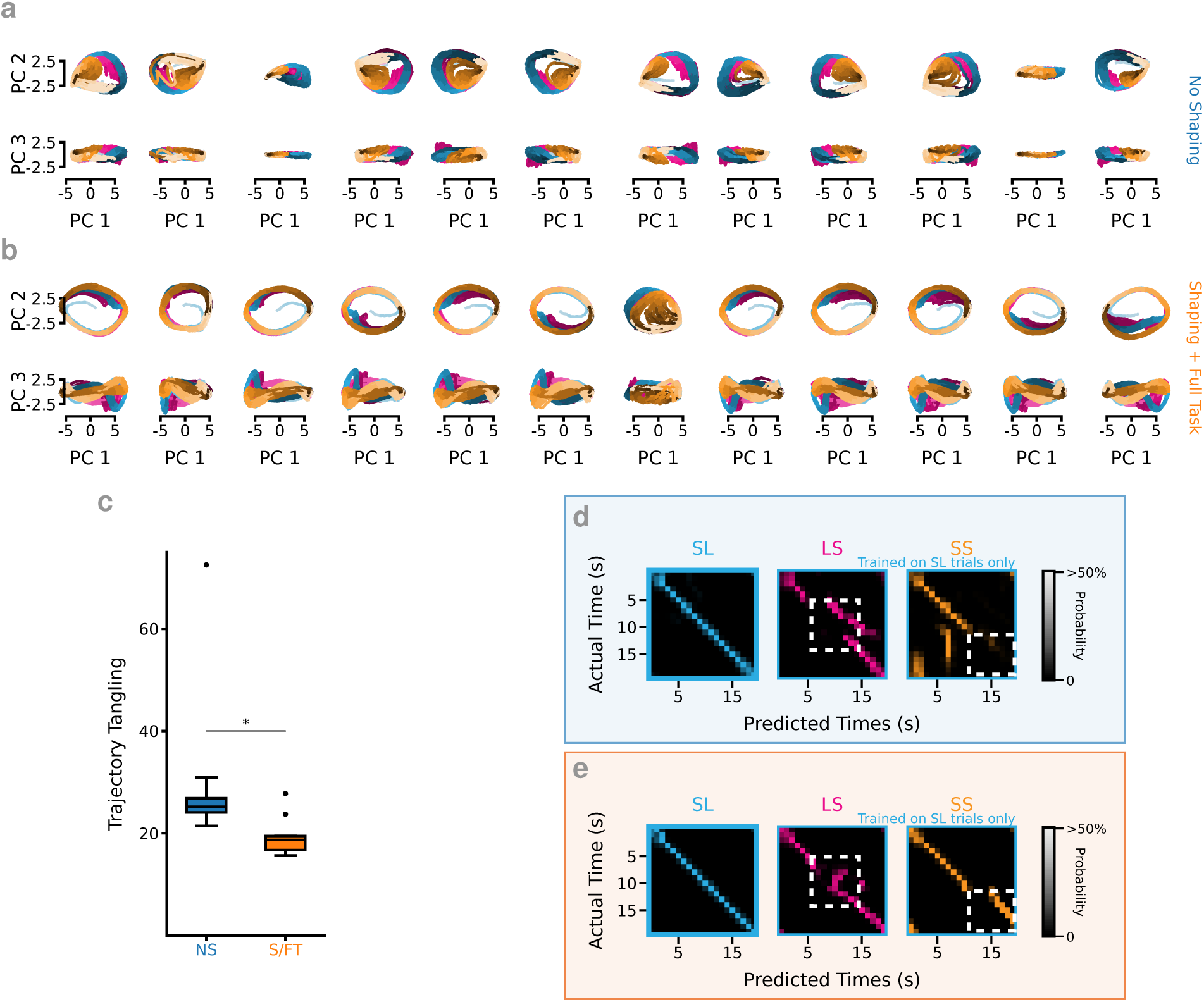
Shaping RNNs constrains network activity to follow smoother state space trajectories. (**a**) PCA state space plots from 12 example NS RNNs independently initialized and trained. *Top*. PC1 and PC2. *Bottom*. PC1 and PC3. (**b**) Same as a but for 12 S/FT RNNs. (**c**) S/FT RNNs show significantly less tangling than NS networks. *n* = 12 NS RNNs, *n* = 12 S/FT RNNs, *p* = 0.02, two sample independent t-test. (**d**) Within- and across-context time decoding for an S/FT RNN from a decoder trained on SL trials. *Left*. LDA based cross-validated decoders trained on SL trials and tested on held out data. *Middle*. The same SL trained decoders tested on LS trials. *Right*. SL trained decoder tested on SS trials. See Figure 2f for LS decoder example. Decoding performed independently and then results averaged across *n* = 10 SL/FT RNNs. (**e**) Same as d but for NS RNN. 2d for LS decoder example.

**Figure S3.**
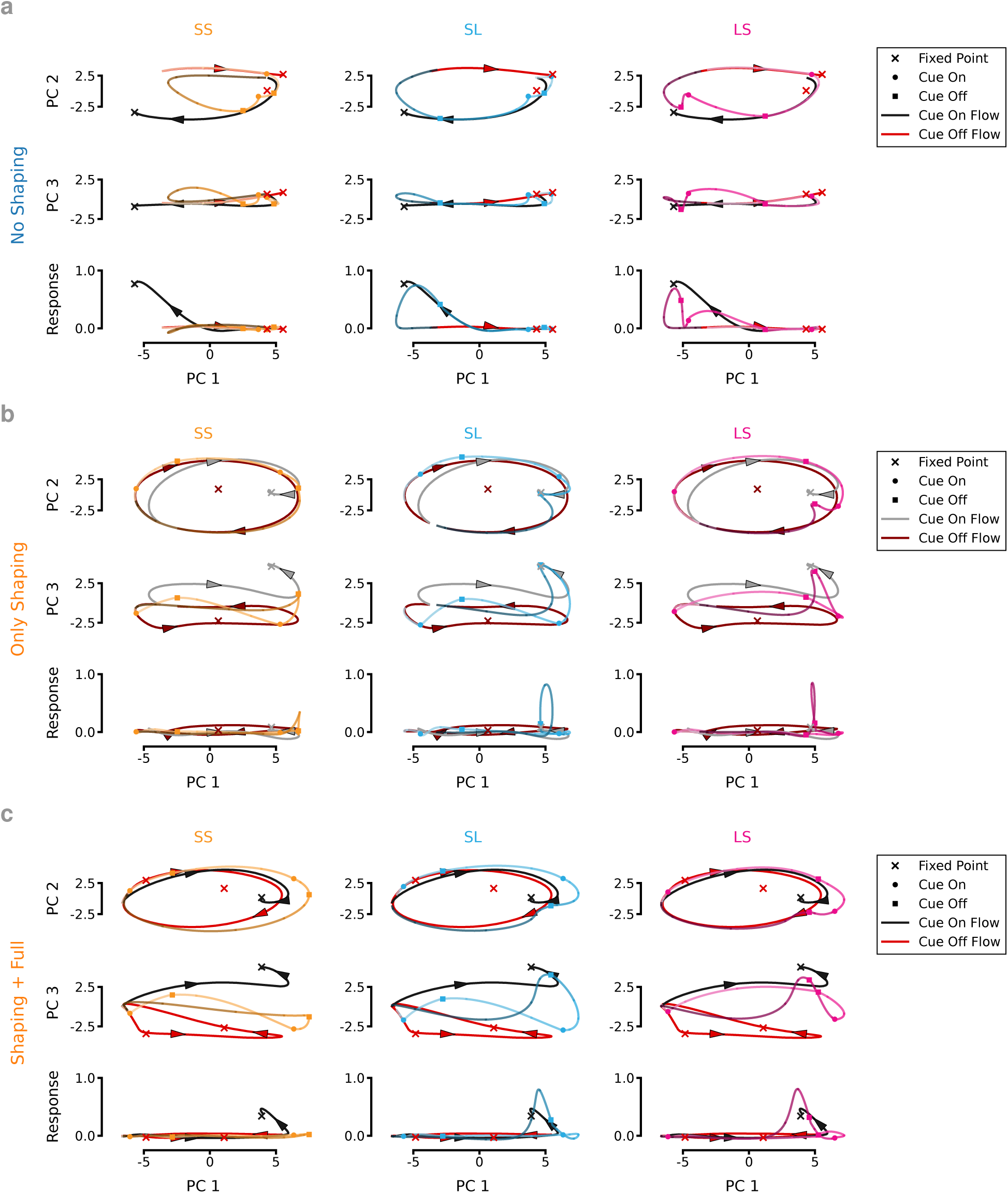
**Dynamical motifs reveal how training approach affects RNNs’ strategy.**(**a**) For each trial type, fixed/slow points for both Cue On and Cue Off modes are shown with state space trajectories superimposed for an NS RNN. *Top*. PC1 vs PC2. *Middle*. PC1 vs PC3. *Bottom*. Response vs PC1. (**b**) Same as a but for an RNN trained on Only Shaping. (**c**) Same as a but for an S/FT RNN.

**Figure S4.**
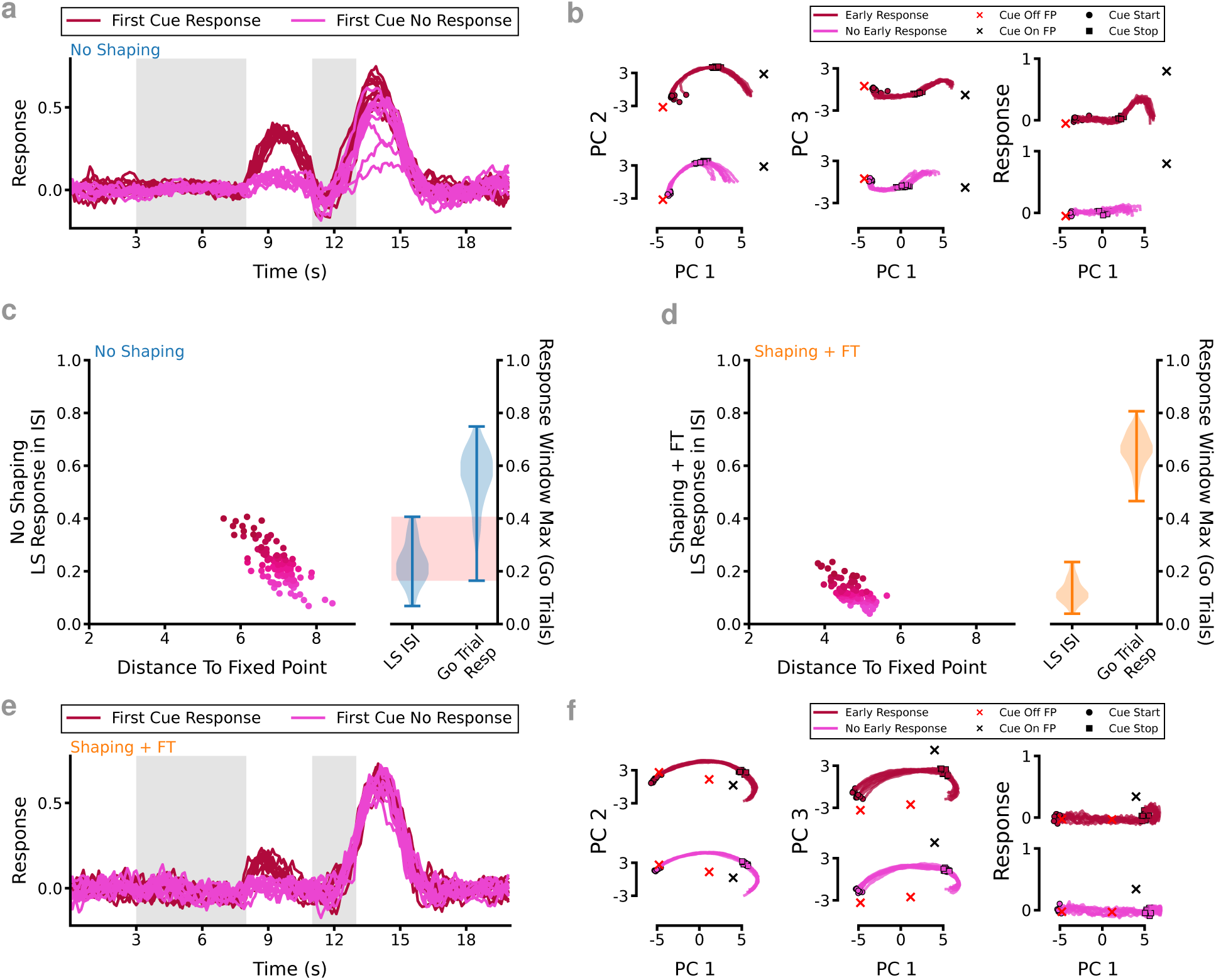
Additional analysis on LS error trials. (**a**) Comparison of response value between the 10% highest response to the first cue on LS trials (early response error) and the lowest for 1 example NS RNN. (**b**) *Top*. PCA state space plots spanning the onset of the first cue until the end of the ISI for 10% largest early response trials. *Bottom*. 10% lowest early response. Note trials where the RNN state is closer to the Cue On fixed point when the cue turns off (square) are more likely to exhibit a response. (**c**) *Left*. Peak response during the ISI compared to euclidean distance to fixed point at the beginning of the ISI shows a strong relationship between how close the RNN is to the fixed point and the likelihood of an early response for *n* = 400 trials in example NS RNN. *Right*. ISI response magnitude peak compared to response window peak magnitude. Overlaps (red) indicates inability to reliably separate “Go” from “No-Go” trials. (**d**) Same as c but for S/FT RNN. Note there is no overlap in the ISI and response window output. (**e**) Same as a but for an example S/FT RNN. (**f**) Same as b but for an example S/FT RNN.

**Figure S5.**
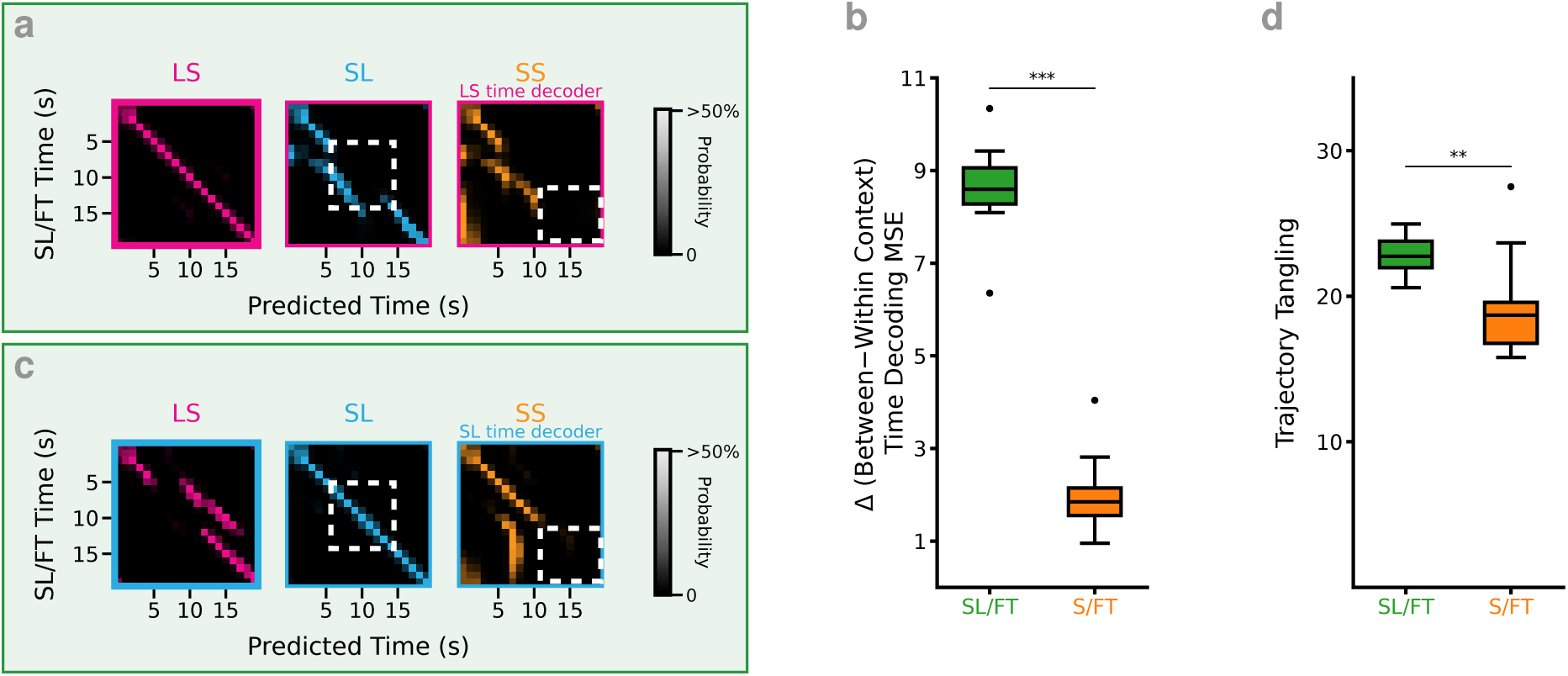
SL/FT networks fail to learn critical abstractions about trial structure. (**a**) Example within- and across-context time decoding for an S/FT RNN. *Left*. LDA based cross-validated decoders trained on LS trials and tested on held out data. *Middle*. The same LS trained decoders tested on SL trials. *Right*. LS trained decoder tested on SS trials. (**b**) Same as a but for decoder trained on SL trials. (**c**) Error comparison for decoding trial time between SL/FT and S/FT RNNs. SL/FT RNNs show decreased performance at cross-context time decoding relative to within, *p* = 2.40 *×* 10*^−^*^13^, *n* = 10 SL/FT RNNs, 12 S/FT RNNs. (**d**) SL/FT RNNs show significantly more tangling than S/FT networks. *n* = 10 SL/FT RNNs, *n* = 12 S/FT RNNs, *p* = 4.62 *×* 10*^−^*^3^, two sample independent t-test.

**Figure S6.**
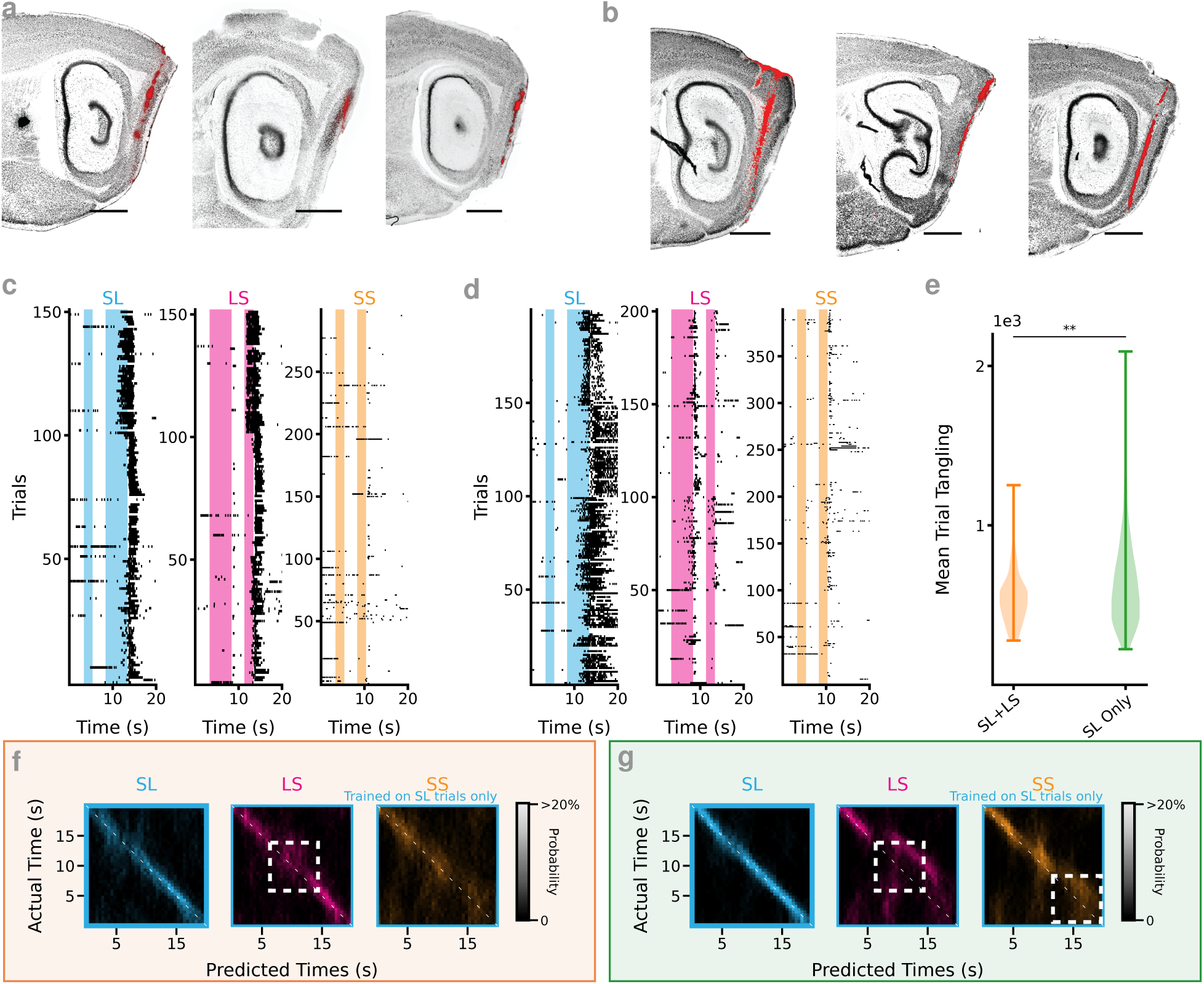
Animals learn shared time-coding axes across trial types—mirroring the dynamics predicted by RNNs. (**a**) Representative section from animal brains trained on S/FT. Nissl stain in black, red dye (DiI/DiD) indicates Neuropixels probe location. Scale bar is 1 mm. (**b**) Same as a but for SL/FT mice. (**c**) Licking activity of all mice trained with SL+LS shaping and then trained on the full task (S/FT mice) on the recording days used for the time decoding analysis. (**d**) Same as c except for the mice trained on SL Only shaping (SL/FT mice). SL/FT mice tend to lick early on LS trials (or not lick at all) resulting in no water reward delivery. (**e**) S/FT shows less tangling than SL/FT (S/FT correct trials only) S/FT *n* = 125 trials from 4 mice, SL/FT *n* = 200 trials from 4 mice, *p* = 1.56 *×* 10*^−^*^3^. Tangling caculated on all recorded units across 25 trials per animal. (**f**) Confusion matrix for time decoding on mice trained with LS+SL shaping (10-fold cross validated). Decoder trained on SL trials then tested on LS, SL, and SS trials. Average of *n* = 5 sessions from 4 mice. See Figure 6g for LS decoder. (**g**) Same as g for SL Only trained animals. Average of *n* = 6 sessions from 3 mice. See Figure 6h for LS decoder.

**Figure S7.**
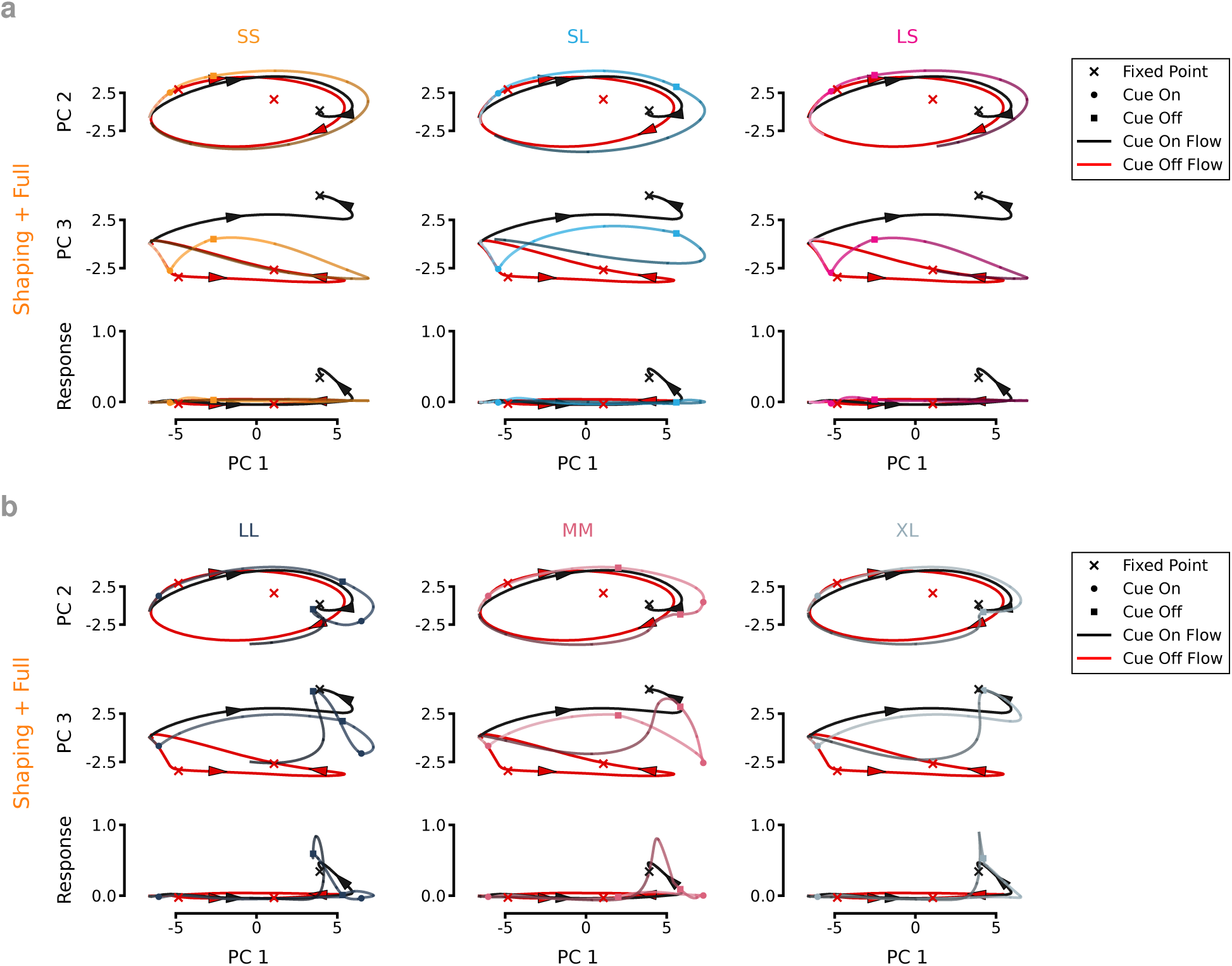
Dynamical motifs make predictions about how RNNs learned strategy extend to novel temporal contexts. (**a**) Both cues contribute to the RNNs decisions to respond. For each trial type, fixed/slow points for both Cue On and Cue Off modes are shown with state space trajectories superimposed for an example S/FT RNN when the first cue is omitted. Note: SS and LS result is effectively the same in this case. *Top*. PC1 vs PC2. *Middle*. PC1 vs PC3. *Bottom*. Response vs PC1. (**b**) The strategy adopted by S/FT RNNs predict a response in reaction to several untrained cue configurations. *Left*. Long-Long (LL) trial type comprised of two 5s cues. *Middle*. Medium-Medium (MM) trial type comprised of two 3.5s cues. *Right*. XL trial type comprised of a single 10s cue.

